# Population model of epigenetic inheritance of acquired adaptation to changing environments

**DOI:** 10.1101/2024.02.16.580565

**Authors:** Dino Osmanović, Yitzhak Rabin, Yoav Soen

## Abstract

Accumulated evidence of transgenerational inheritance of epigenetic and symbiotic changes begs the question of under which conditions inheritance of acquired changes can confer long-term advantage to the population. To address this question, we introduce a population epigenetics model of individuals undergoing stochastic and/or induced changes that are transmitted to the offspring. Potentially adaptive and maladaptive responses are represented, respectively, by environmentally driven changes that reduce and increase the individuals’ rate of death (i.e. reduction and increase of selective pressure). Analytic solution in a simplified case of exposure to two types of dynamic environments shows that inheritance of changes that transiently alleviate the selective pressure confers long-term advantage even when the transmitted state is maladaptive to the offspring. The benefits of inheriting environmentally driven changes that reduce the death rate within a lifetime include escape from extinction under a wide range of conditions. These advantages are even more pronounced in populations with imperfect inheritance and/or age-dependent decline in fertility. These findings show that inheritance of non-genetic changes can have tremendous benefits for the population on timescales that are much longer than the lifetime of an individual.

## I. INTRODUCTION

Non-genetic inheritance of phenotypic changes that are acquired during the lifetime of individuals has been reported in both animals and plants [1–5], including in response to environmental stressors/cues (heat stress, endocrine disruptors, viral infection, nutrient limitations and diet composition, toxic exposure, traumatic experiences, etc.) [6–21]. Studies of diverse cases have also identified distinct mechanisms that can either mediate or support inheritance of acquired changes, such as: resistance to viral infection by non-coding RNAs in C. elegans [13, 14], DNA methylation-based persistence of endocrine disruption impacts in rats [8, 14], involvement of histone modifications in trasgenerational persistence of responses to temperature in Drosophila [6, 22, 23], inheritance of piRNA-dependent RNA silencing in C. elegans by small RNA and chromatin components in the nucleus [24], inheritance of DNA methylation changes in heat exposed wild guinea pigs [25] and in fish that are exposed to hypoxia [26] and hydrogen sulfide–rich environments [17], inheritance of toxin-induced phenotypes in Drosophila by altered deposition of maternal RNAs [27] as well as by persistence of changes in the symbiotic gut microbiota [28, 29]. Transgenerational impacts may involve stochastic and/or deterministic changes and can be beneficial or non-beneficial [30]. A hallmark example of adaptive changes that are acquired de novo within a generation and epigenetically inherited for dozens of generations is provided by the acquisition of immunity to new viruses in C. elegans [9]. This immunity is acquired by a dedicated machinery for using the virus to mount a specific defense in the form of small RNA which is subsequently amplified and transmitted across generations of offspring. Transgenerational persistence of harmful impacts has also been demonstrated, such as inheritance of metabolic dysregulation and increased susceptibility for obesity in response to environmental toxicants and altered nutrition [15, 31].

While the inheritance of non-genetic variations can clearly affect the conditions of selection acting on a population [32–35], the impacts on the population depend on the magnitude, rate and type of environmental change [36–38], host-symbiont interactions [39, 40], as well as population structure (manifested, for example, by kin phenotypic correlations [41]. This gives rise to a complex repertoire of possible outcomes, including scenarios in which the impact of epigenetic inheritance of a specific phenotype turns from being beneficial to non-beneficial and vice versa [30, 36, 42]. Due to the complexity of possible scenarios, it is not clear whether, and under which conditions, the inheritance of acquired variations confers long-term benefit to the population [35, 43]. Addressing this question requires a modelling framework capable of drawing conclusions that are applicable to a wide range of populations and environmental regimes. An analogous problem of assessing impacts of genetic and/or phenotypic variations in the populations under complex, time-varying ecology is addressed by combining population genetics with population ecology [39, 40, 44–54]. While enabling consideration of ecology-driven changes in population structure on timescales that are smaller than one generation [49, 50, 55], the traditional models do not account for inheritance of changes that are rapidly acquired (or made) by individuals. Initial evaluation of adaptive contributions of epigenetic inheritance under changing environments considered the transmission of epigenetic states that are assumed to be advantageous to the offspring [56, 57]. In general, however, epigenetic variations are not necessarily advantageous. Moreover, since parents and offspring may be exposed to different environments, the transmission of changes that were beneficial to the parents may not be beneficial (and can even be detrimental) to the offspring. Comprehensive analysis of the effects of inheriting acquired changes therefore requires a modelling framework that accommodates for different scenarios and links changes in individuals to longer-term impacts on the population. Towards this goal, we introduce a population model of individuals undergoing dynamic changes that are either partially or fully transmitted to the offspring. The model formulation is general enough to consider any type of stochastic or environmentally induced changes that take place during the lifetime of individuals and can persist in their offspring. This includes internal changes in individuals (e.g. epigenetic and symbiotic variations) as well as niche construction changes that they make in their environment. The acquired changes include stochastic variations as well as directed responses to the environment that either reduce or increase the individuals’ rate of death (thus reducing or increasing the selective pressure).

Model formulation and analysis are described in the following order: In section II we extend the traditional framework by formulating a model that considers inheritance of stochastic and environmentally-induced changes that are acquired during the lifetime of individuals. In section III we derive analytic solution for specific types of populations, and determine the resulting impacts on population size in a stationary environment as a function of population and environmental parameters. In sections IV to VI, we determine how the inheritance of acquired changes influences population size and sustainability under two distinct types of dynamic environments. Section VII demonstrates how the analytic solution can be extended to scenarios of imperfect inheritance and age-dependent decline in fertility. Section VIII discusses the findings and limitations of this modeling framework.

## II. MODEL FORMULATION

We consider an evolving population of *N* individuals, wherein the rates of birth and death of the *i*^*′*^*th* individual depend on its age *T*_*i*_(*t*) and state variable *χ*_*i*_(*t*) at time *t*. This state is taken to represent any type of variable, phenotype, or process that can be acquired and/or modified by individuals during their lifetime, and subsequently transmitted with some fidelity to the offspring. This includes epigenetic and symbiotic variables as well as elements in the environment that are created and/or affected by the individuals (niche construction).

To derive a model for the time evolution of such a population, we extend the McKendrick–Von Foerster equation [50, 58–60] to include stochastic and non-stochastic changes in *χ*_*i*_ that take place during a lifetime. The distribution of ages *T* and states *χ* in the population at time *t* is given by the sum over the individuals:

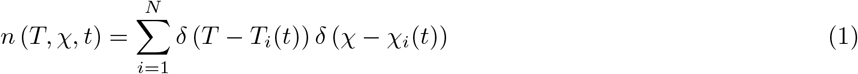

where *δ*() is the Dirac delta function. Note that the age and state variables of the distribution, *T* and *χ*, are independent parameters that should not be confused with actual ages *T*_*i*_(*t*) and states *χ*_*i*_(*t*) of specific individuals at time *t*. The population size *N* (*t*) is, in turn, obtained by integrating *n* (*T, χ, t*) over *T* and *χ*. To derive a differential equation that describes the time evolution of the population distribution, we consider age- and state-dependent rates of death and replication (birth) *P*_*D*_(*T, χ*) and *P*_*R*_(*T, χ*), respectively and take the temporal changes in existing individuals into account by assigning suitable dynamical rules for the ages *T*_*i*_(*t*) and states *χ*_*i*_(*t*) of individuals. Since age increases linearly with the passage of time, the time evolution of *T*_*i*_(*t*) is trivial:

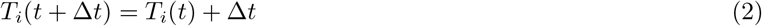

For a sufficiently small time duration Δ*t*, the time evolution of *χ*_*i*_(*t*) can be generally represented as a combination of directed and non-directed changes, governed by a drift function *f* (*χ*_*i*_, *t*) (assumed to be a function of the state *χ*_*i*_ and of time *t* but not of the age *T*_*i*_), and a Wiener-type stochastic process, *ξ*(*t*):

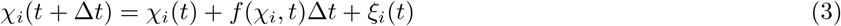

Assuming that *ξ*_*i*_(*t*′) is a random Gaussian distributed variable, its second moment can be expressed as ⟨*ξ*_*i*_(*t*)*ξ*_*j*_(*t*^*′*^)⟩ = 2*Dδ*_*ij*_*δ*(*t* − *t*) where *D* plays the role of a ‘diffusion coefficient’ in *χ* space. To shift from an individual-level to a mean field description at the population-level, we further assume that the stochastic variable *ξ* is independent of *χ* and that the function *f* and the statistical properties (e.g., variance) of *ξ* are the same for all individuals in the population.

Taken together with eq. 1, the time evolution of the population distribution, *n* (*T, χ, t*), is specified by a differential equation with an average rate of death, *P*_*D*_(*T, χ, t*), diffusion and drift (eq. 3), subject to the boundary conditions that age is bounded from above, and that the distribution of newborns, *n* (*T* = 0, *χ, t*), is governed by the rate of replication, *P*_*R*_(*T, χ, t*), and the inheritance function’, *I*(*χ* − *χ*^*′*^):

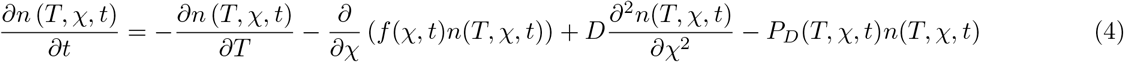

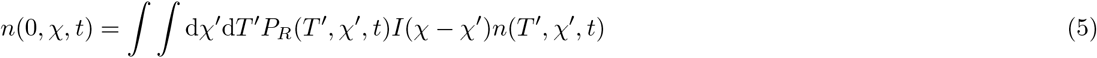

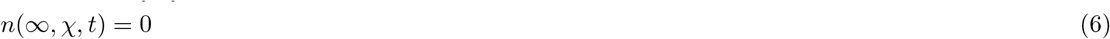

The derivation of eq. 4 is provided in Supplementary Information (SI), section I. The effects of directed and stochastic changes in state are represented in eq. 4 by the drift and diffusion terms (second and third terms respectively, on the right-hand-side). The other two equations correspond to boundary conditions at ages *T* = 0 and *T* = ∞, respectively. The fidelity of transmission of states across generations is, in turn, specified by the inheritance function *I*(*χ*−*χ*^*′*^) in eq. 5. Perfect inheritance corresponds to *I*(*χ*−*χ*^*′*^) = *δ*(*χ*−*χ*^*′*^), and imperfect inheritance can be conveniently represented by a Gaussian function whose width determines the extent of deviation from perfect inheritance. The consideration of inheritance function that depends only on the distance of *χ* from *χ*^*′*^ is a crude representation of the complexity of replication, but it is nonetheless useful for recapitulating similarities between populations of offspring and those of their parents. Equation 6 is a boundary condition which states that the population contains no individuals of infinite age.

To model acquisition of changes that either reduce or increase the selective pressure on timescales that can be shorter than one generation, we consider a drift function *f* (*χ, t*) that is proportional to the derivative of *P*_*D*_(*χ, t*) with respect to *χ*, namely:

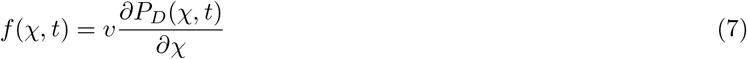

where *v* is a drift coefficient controlling the rate at which the change of *χ* increases or decreases the death rate. A drift with negative *v* pushes *χ* towards states of lower death rate, representing a scenario in which individuals respond to selective pressure by changing their *χ*_*i*_ values along a direction that reduces the pressure. A positive *v*, on the other hand, corresponds to changes in a direction that increases the selective pressure. Such a drift function appears naturally in population dynamics models of slowly breeding host organisms that live in symbiosis with fast breeding bacteria (see SI, section II). Note that consistent with our assumption that *f* is a function of *χ* and *t* only, in the following we will assume that the death rate depends only on the state parameter *χ* and on time *t*, but not on the age parameter *T*.

## III. TIME EVOLUTION OF THE POPULATION IN STATIONARY ENVIRONMENT

To investigate the time evolution of a population in a stationary (time-independent) environment, we consider a death rate that depends quadratically on *χ* and linearly on population size, *N* (*t*) = ∬ d*χ*^*′*^d*T* ^*′*^*n*(*T* ^*′*^, *χ*^*′*^, *t*) (as in logistic growth):

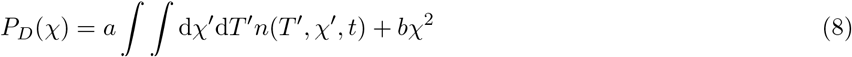

The quadratic term represents a plausible approximation of the effect of environmental selection on the variable *χ*, at the vicinity of the value that minimizes the death rate.

To analyze the population dynamics under this form of death rate, we start by considering the case of constant proliferation rate, *P*_*R*_(*T, χ*) = *r*, and perfect inheritance *I*(*χ*−*χ*^*′*^) = *δ*(*χ*−*χ*^*′*^). In this case we have three population-specific parameters (drift *v*, diffusion *D* and replication rate *r*) and two environmental parameters, *a* and *b*, determining, respectively, the strength of resource limitation (‘carrying capacity’) and the strength of the selective pressure on *χ*. Due to the carrying capacity term (proportional to *N* (*t*)) in eq. 8, the time evolution of the distribution, eq. 4, includes a term that is quadratically dependent on *n*(*T, χ, t*). We nonetheless find that this case has an analytic solution that can also be extended to populations of individuals with age-dependent decline in fertility and imperfect inheritance. The solution is obtained by subjecting eq. 4 to a Laplace transform with respect to *T* and a Fourier transform with respect to *χ*, and can be conveniently expressed in terms of the integral transform of the population distribution (the characteristic function 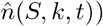:

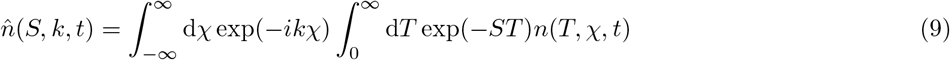

Note that the population size at time *t* is given by the zero modes (S=0,k=0) of the characteristic function, i.e. 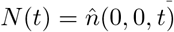. As detailed in section III of the SI, the time evolution of the characteristic function for a population with constant fertility and perfect inheritance is fully specified by the *S* = 0 mode of the characteristic function, 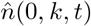:

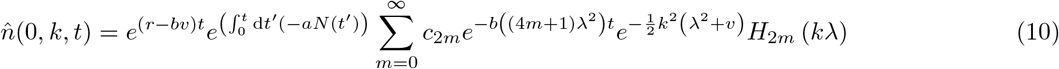

where we defined

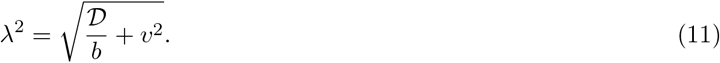

The coefficients *c*_2*m*_ depend on the choice of initial conditions, and the functions *H*_2*m*_ are the Hermite polynomials. Various other statistical properties of the population (e.g., the average age) are defined by derivatives of 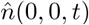. The differential equation governing the time evolution of the population size is obtained by combining eq. 10 with the integral transforms of eq. 4 (see SI section III for complete derivation):

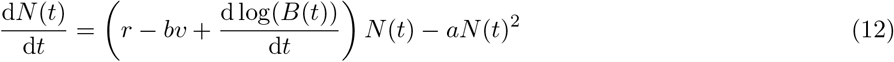

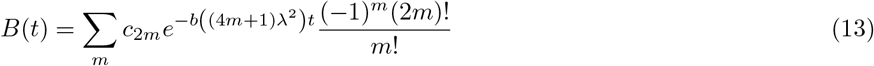

Note that eq. 12 can be viewed as logistic growth with a time-dependent growth rate *r* −*bv* + d log(*B*(*t*))*/*d*t*.

Some of the generic effects of the population parameters *v, D* and *r* and the environmental parameters *a* and *b* are revealed by the equilibrium distribution to which the population relaxes in the long time limit (starting from arbitrary initial conditions), 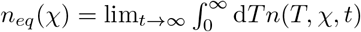, and the equilibrium size of the population, *Neq*:

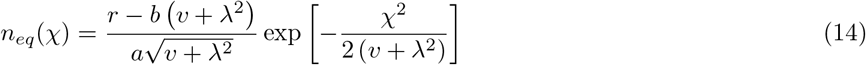

and

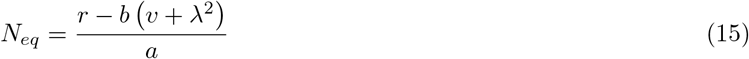

The age-dependent equilibrium distribution decays exponentially with *T*, i.e. *n*_*eq*_(*T, χ*) = *rn*_*eq*_(*χ*) exp(−*RT*). The derivation of the above expressions is given in section IV of the SI. The dependence of total number of individuals *N*_*eq*_ and of the distribution *n*_*eq*_(*χ*) in equilibrium, on the parameters *v* and *D* is summarized in figures 1*A* and *B* respectively:

**FIG. 1:**
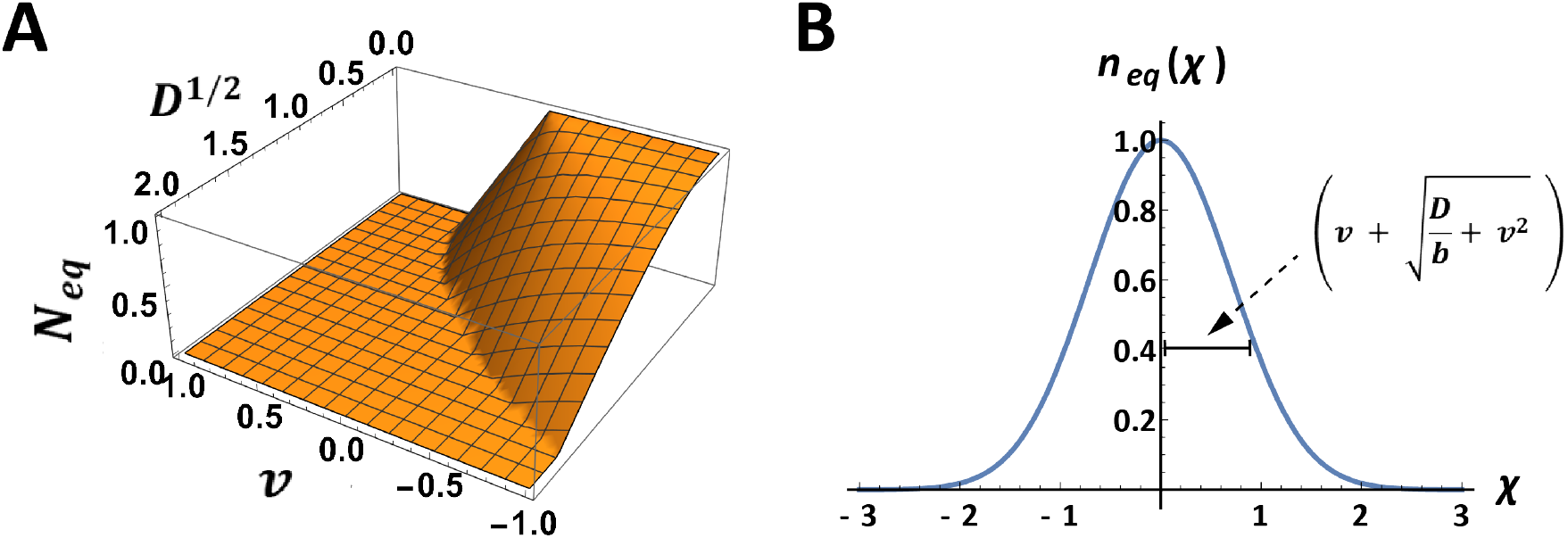
Properties of the equilibrium state. A) A 3d plot of the total amount of individuals *Neq* as a function of the drift parameter *v* and the diffusive parameter *D*^1*/*2^ (with *r* = *a* = *b* = 1). B) The equilibrium distribution *neq* (*χ*) is given by a Gaussian function with a variance that depends on *D* and *v*.

Equation 15 shows that *N*_*eq*_ decreases with *D* and increases with more negative *v*. Despite this benefit of more negative *v*, the equilibrium size of a population with constant *r* in a stationary environment is maximal when both *D* and *v* are zero (i.e. when *D* = 0, faster reduction of the selective pressure within a generation cannot increase the steady state size of the population beyond the that of a population with *v* = 0). This can be understood by realizing that in the limit of *D* → 0 and *v* → 0, the equilibrium distribution *n*_*eq*_(*χ*) becomes proportional to a delta function at a value of *χ* that minimizes the selective pressure (adaptive peak). A non-vanishing *v* broadens this distribution towards states of increased rate of death, resulting in lower *N*_*eq*_. However, in any realistic scenario, *D* is larger than zero and *N*_*eq*_ increases as a function of the capacity to reduce the selective pressure within generation (negative *v*). Consideration of changes in more than one parameter shows that the same population *N*_*eq*_ can be maintained by different combinations of the parameters *v, D* and *r* (Fig. 1A). For example, a given increase in *N*_*eq*_ due to reduction of selective pressure within generation (negative *v*) can also be achieved by faster replication (larger *r*) in populations of individuals that are incapable of reducing the selective pressure (i.e. having *v* = 0). Similarly, the negative impact of a larger *D* can be counteracted either by faster replication (larger *r*) or by faster reduction of the selective pressure (more negative *v*). Thus, as long as the environment is constant, the same population size can be achieved by distinct ‘strategies’, corresponding, respectively, to altering the rate of replication and acquiring heritable changes that modify the selective pressure, either by directional changes (via *v*) or by stochastic variations (*D*).

## IV. CONSIDERATION OF DYNAMIC ENVIRONMENTS

To investigate implications of inheritance of acquired changes in a dynamic (changing with time) environment, we consider shifts in the state (*χ*_*min*_) that minimizes the death rate. For example, a sudden shift from *χ*_*min*_ = 0 in an old environment to *χ*_0_ in a new environment is defined by changing the death rate in eq. 8 to:

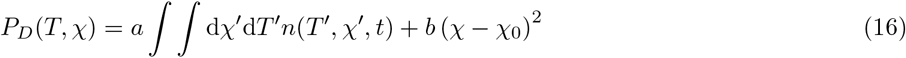

Prior to this sudden change, the rates of death and birth were equal for a population at equilibrium. The change in *χ*_*min*_ increases the death rate and drives dynamic changes in the distribution *n* (*T, χ, t*) until the population reaches a new equilibrium. Since the rate of death *P*_*D*_(*χ*) and the resulting drift *f* (*χ*) at equilibrium around *χ* = 0 in the old environment are the same as those around *χ*_0_ in the new environment, *N*_*eq*_ in the new environment is the same as in eq. 15 and the new equilibrium distribution *n*_*eq*_ is obtained by replacing *χ* in eq. 14 with *χ*−*χ*_0_. Parameter-dependent analysis of populations that were at equilibrium in the old environment (section V in SI), reveals that the characteristic time for relaxation in the new environment (in units of 1*/r*) decreases with negative *v* (Fig. 2A), indicating that relaxation is expedited by the ability to inherit acquired changes that reduce the selective pressure within generation (*v <* 0). On the other hand, the qualitative impact of stochastic changes (*D*) as well as changes that increase the selective pressure (*v >* 0), depend on the magnitude of these changes (Fig. 2A). If both *v* and *D* are sufficiently small, a larger *D*, or alternatively more positive *v*, contributes to expedited adaptation by broadening the width of the initial distribution, thus increasing the overlap between the tail of the initial distribution and the center of the new one. Beyond certain levels of *v* and *D*, however, further increase in positive *v* or in *D* has the inverse effect of increasing the relaxation time, which diverges at finite values of positive *v* and *D* (Fig. 2A).

**FIG. 2:**
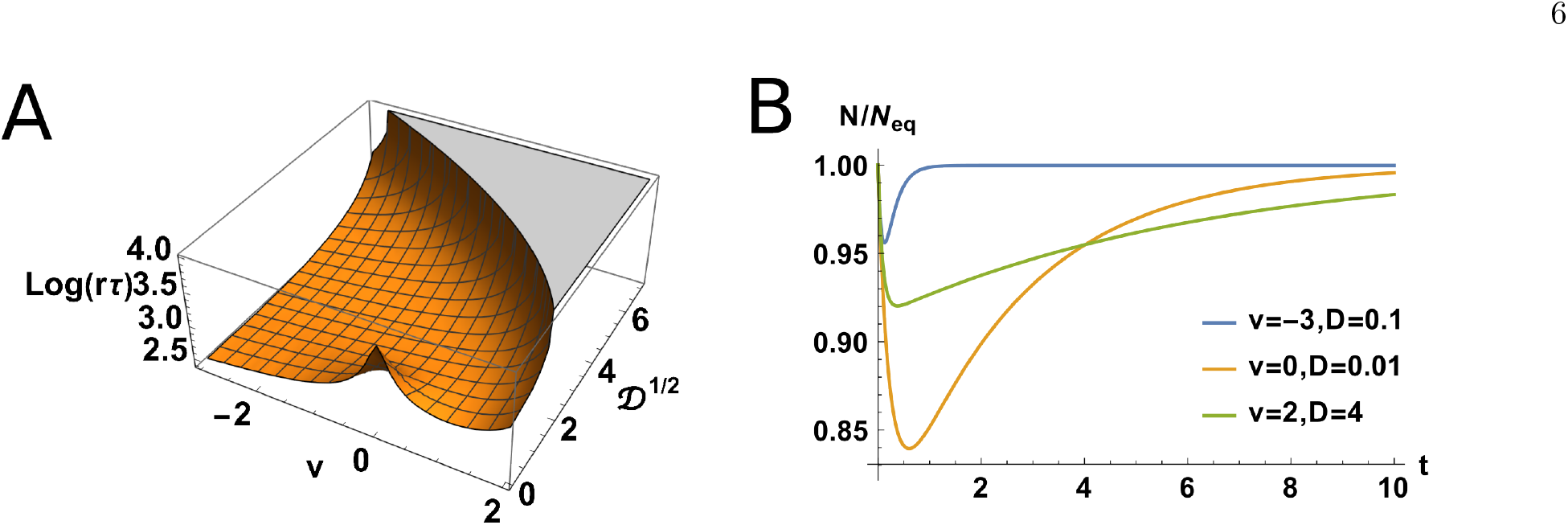
Shifting environments. A) Following an environmental shift, the population is perturbed and relaxes to a new steady state. Displayed is the characteristic relaxation time *τ* (in units of 1*/r*) of populations of different *v* and *D* with *r* = 5 after a shift Δ*χ* = 1. The relaxation time (defined as the time it takes the population to recover), diverges in certain regions of the parameter space *v, D*. B) Examples of relaxation for 3 different sets of parameters. All examples display fast initial decrease followed by slower recovery.

In the case of a single transition between stationary environments, the benefits of inheriting acquired changes that reduce the selective pressure are fully expected. However, if the environment continues to change, the inherited modifications in *χ* may be outdated and no longer advantageous. We therefore sought to investigate the potential long-term impacts of inheriting acquired changes for different scenarios of change in the environment. For that, we considered two different types of dynamical change: (a) periodic cycles of jumps in the selective pressure (Type I dynamics), and (b) unidirectional jumps in the selective pressure (Type II dynamics). We incorporate these changes into the model by assigning *χ*_0_ with dynamical rules corresponding to the two types of change. Periodic jumps between states *χ*_0_ and −*χ*_0_ (Type I dynamics) with Δ*χ* = 2*χ*_0_ and period Δ*t* are specified by:

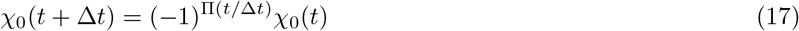

where Π (*t/*Δ*t*) is a square wave function oscillating between 0 and 1 with period Δ*t*. Unidirectional jumps (Type II dynamics) at a rate Δ*χ/*Δ*t* are, in turn, specified by:

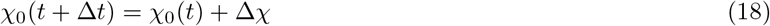

In both cases, the magnitude and rate of change are determined by the additional environmental parameters Δ*χ* and Δ*t*. The calculation of the final distributions for both types of dynamics is outlined in section VI of the SI.

## V. CONDITION FOR POPULATION SURVIVAL

≥

To derive an approximate condition for survival under these changes in the environment, we analyzed the dynamics of the slowest mode in the expansion of the population distribution. Realizing that the population survives only if the slowest mode is fully recovered before the next environmental change (i.e. *N* (Δ*t*)≥*N* (0)) leads to a necessary condition for survival (full derivation is provided in section VII of the SI):

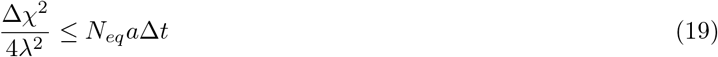

Substituting for *N*_*eq*_ in eq. 15 gives:

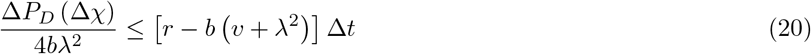

where Δ*P*_*D*_ (Δ*χ*) = *b*Δ*χ*^2^ corresponds to the initial jump in the rate of death due to the sudden displacement by Δ*χ* from the state that minimized the death rate prior to the jump. Recalling that *r*Δ*t* is the number of births within a time interval Δ*t* yields:

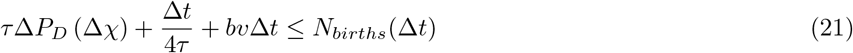

where *τ* = 1*/*4*bλ*^2^ is the characteristic timescale for broadening of the distribution *n*(*χ*). To achieve complete recovery within Δ*t*, the number of births must compensate for the number of deaths in this time interval. The latter is displayed on the left-hand-side of eq. 21 as a sum of 3 components (left to right): (i) number of “excess” deaths (due to the shift by Δ*χ*_0_ during the characteristic time for alleviating the increase in death rate by broadening the distribution toward the new minimum, (ii) number of deaths in time Δ*t* due to limited carrying capacity, and (iii) the reduction (for *v <* 0) or increase (for *v >* 0) in the number of deaths by the directed change (drift) in *χ* over time Δ*t*.

Note that the above analysis neglects the contributions of higher modes and therefore provides only an approximation to the exact necessary conditions for survival. Since higher modes decay faster than the lowest mode, the lowest mode approximation is expected to be better for higher values of Δ*t*. This is clearly observed in figures 3C and 4B. The approximation clearly breaks down for small Δ*t* in the case periodic switching (figure 3C), concurring with the expectation that for sufficiently rapid periodic changes of the environment, the population can survive even without adapting to the higher death rate in the new environment.

**FIG. 3:**
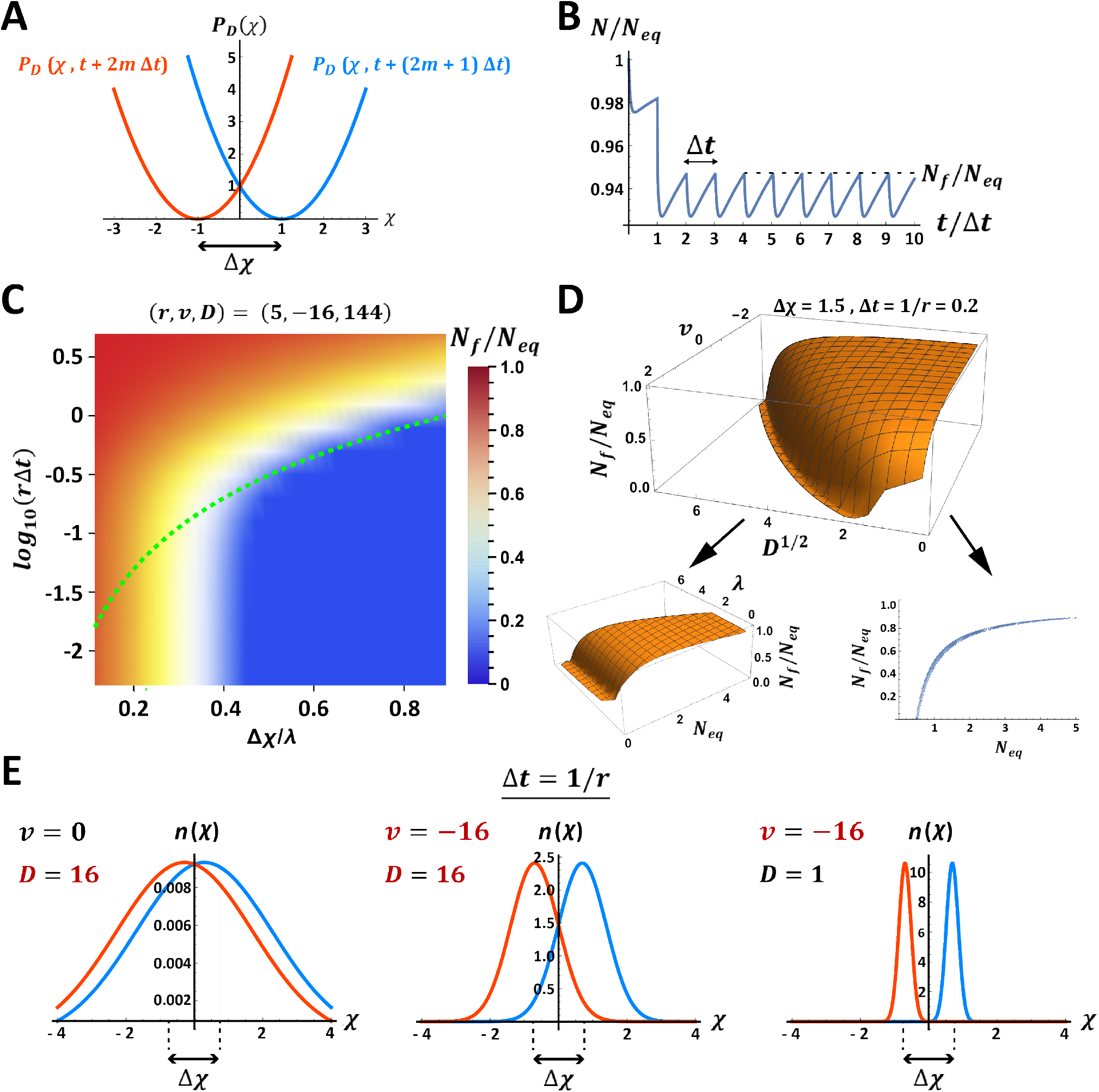
Population outcomes under periodic shifts of the environments (Type I dynamics). A) Illustration of periodic switching of a parabolic death rate function, corresponding to back-and-forth jumps (of magnitude Δ*χ*=2) in the adaptive peak, with a duration of Δ*t* between consecutive jumps. B) Example of population recovery following a transition from a stationary environment to the periodically switching environment shown in (A). C) Normalized maximal level of the recovered population, *N*_*f*_ */N*_*eq*_, as a function of *r*Δ*t* and 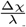 for a specific choice of *r, v*, and *D*. Note the continuous transition from population survival to extinction in (Δ*χ*, Δ*t*) space. Green dashed line displays the boundary of the necessary condition for survival, Eq. 19 (below this line, the population necessarily goes extinct). D) Landscape of *N*_*f*_ */N*_*eq*_ as a function of *v* and *D*^1*/*2^ for a given magnitude of the environmental switch (Δ*χ* = 1.5) and duration between switches (Δ*t* = 0.1). Survival is supported by more negative values of *v* and smaller values of *D*. Lower panels: *N*_*f*_ */N*_*eq*_ vs. *N*_*eq*_ (right), and*N*_*f*_ */N*_*eq*_ as a function of *N*_*eq*_ and *λ* (left). E) Effect of negative *v* on the shape of the distribution *n*(*χ*) immediately prior to each switch. Shown for *v* = −16 with high and low *D* (middle and right panels) vs. *v* = 0 and high *D* (left).

## VI. LONG-TERM IMPACTS OF THE INHERITANCE OF ACQUIRED CHANGES IN DYNAMIC ENVIRONMENTS

To simplify the investigation of the effects of *v, D* and *r* in changing environments, we eliminated the dependence of the initial equilibrium population on these parameters by measuring the population size *N* (*t*) in units of *N*_*eq*_ prior to changes in the environment. We start by considering the case of periodically switching environment (Type I dynamics-see Fig. 3A). Example of a typical trajectory of *N* (*t*)*/N*_*eq*_ for a surviving population in a switching environment is shown in Fig. 3B. Each switch triggers a population decline due to the sudden increase in selection pressure followed by potential recovery that is mediated by a gradual shift of the distribution toward states of lower selective pressure. This process converges to periodic saw-like pattern of population size bounded by constant levels of minimal and maximal population size. The maximal level of the recovered population, *N*_*f*_, is a measure of how resilient the population is to the environmental perturbation. Analysis of *N*_*f*_ */N*_*eq*_ as a function of Δ*χ* and Δ*t*, reveals a sharp separation between survival and extinction (Fig. 3C). A similar separation is noted by plotting *N*_*f*_ */N*_*eq*_ as a function of *v* and *D*^1*/*2^ for a specific choice of Δ*χ* and Δ*t* (Fig. 3D), demonstrating that the recovered population decreases with larger *D* and increases with more negative *v*. Since the magnitude of the drift is proportional to both *v* and *χ*, negative *v* confers two benefits: shifting the distribution towards the adaptive peak and narrowing it around the peak (Fig. 3E). A sufficiently negative *v* also enables sustainability of populations that would otherwise go extinct. Plotting *N*_*f*_ */N*_*eq*_ vs. *N*_*eq*_ and *λ* at fixed Δ*χ* and Δ*t* (Fig. 3D, bottom left panel), shows that *N*_*f*_ is fully determined by the initial equilibrium size of the population, *N*_*eq*_ and by the value of *λ*. For a sufficiently small Δ*t, N*_*f*_ is largely determined by the initial equilibrium size, *N*_*eq*_ and is only weakly dependent on *λ* (Fig. 3D, bottom right and left panels). Below a critical level of *N*_*eq*_, the population goes extinct and immediately above this cutoff, *N*_*f*_ */N*_*eq*_ scales as 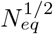 (not shown), curiously reminiscent of critical behavior in Ising-like systems [61].

In the case of Type II dynamics (Fig. 4A), the state that minimizes the selective pressure, *χ*_0_(*t*), changes at a rate Δ*χ/*Δ*t*, thus presenting an increasing challenge to the population. Nonetheless, for given Δ*χ* and Δ*t*, there exists a range of parameters *v, D, r, a* and *b*, supporting a non-vanishing value of *N*_*f*_. For the most part, the effects of environmental (Δ*χ*, Δ*t*) and population parameters (*v, D*) on *N*_*f*_ */N*_*eq*_ under unidirectional challenge and under periodical switching of the environment, are qualitatively similar (compare Figs. 4B and Figs. 3C). However, in contrast to exposure to periodically switching environment, in the case of unidirectional challenge, populations with small enough *D* may benefit from slow acquisition of changes that increase the selective pressure. This is demonstrated by the observed increase of *N*_*f*_ */N*_*eq*_ with *v* for small enough levels of positive *v* (Figs. 4C). This benefit stems from the contribution of positive *v* to the broadening of the distribution, enabling survival of individuals with a state that is sufficiently close to *χ*_0_(*t*).

**FIG. 4:**
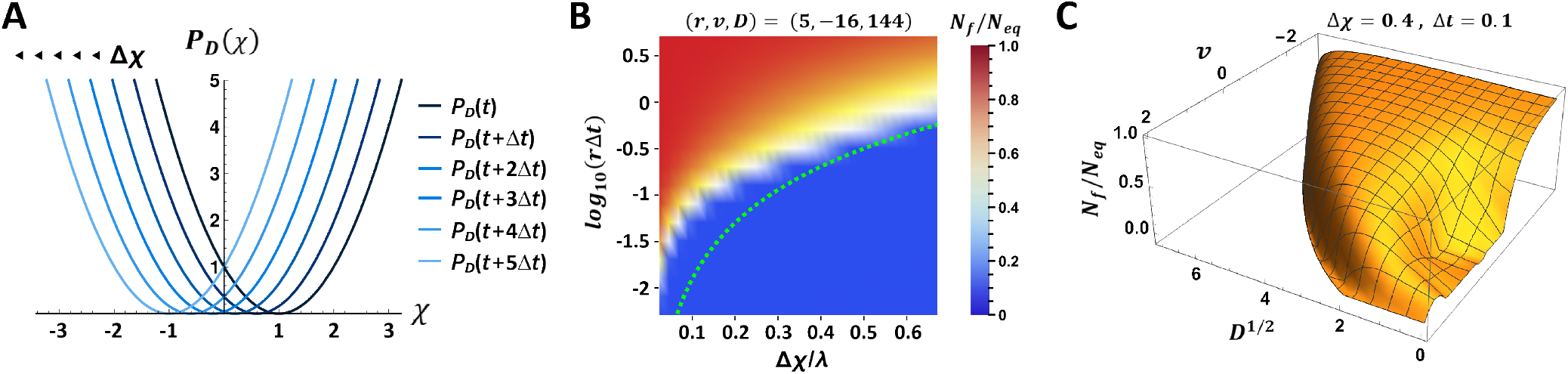
Population outcomes under successive shifts of the environment (Type II dynamics). A) Unidirectional jumps of the adaptive peak by Δ*χ* every Δ*t*. B) *N*_*f*_ */N*_*eq*_ as a function of *r*Δ*t* and 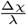 for the same choice of *r, v*, and *D* as in Fig. 3C. Green dash line displays the boundary of the necessary condition for survival, Eq. 19. C) Landscape of *N*_*f*_ */N*_*eq*_ as a function of *v* and *D*^1*/*2^ for unidirectional jumps of magnitude Δ*χ* = 0.4 and Δ*t* = 0.1 between jumps. The approximated boundary of the region below which the population necessarily goes extinct is indicated by the green dash line. While negative *v* is generally beneficial (as in the case of periodic switching), the effects of positive *v* and *D* change from a positive contribution to population survival at small enough *v* and *D* to a deleterious impact at larger values of *v* and *D* (in contrast to the case of periodic switching; figure 3).

## VII. IMPERFECT INHERITANCE AND AGE-DEPENDENT DECLINE IN FERTILITY INCREASE THE IMPACT OF ACQUIRED CHANGES

To investigate how the long-term impacts of inheriting acquired changes are affected by age-dependent fertility and fidelity of inheritance, we consider a population of individuals whose rate of reproduction declines with age and whose state is imperfectly transmitted to the next generation. To maintain the ability to derive analytic solutions despite these additional complications, we assume that the replication rate declines exponentially with age

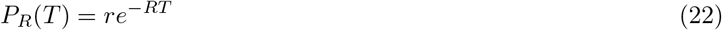

where *R* is the decay rate of fertility with age. We would like to stress that the exponential form of the replication rate, eq. 22, is chosen to simplify the analytical analysis rather than to represent biological reality. While it is clearly non-realistic to reach maximal reproduction rate at age zero, consideration of more realistic forms of age-dependent replication rates leads to similar conclusions (SI, section X). As shown in the supplementary information (SI, section VIII), the decline in fertility allows us to decouple the age at which the individuals are most fertile from the average age at reproduction, 1*/R*. In addition, the average lifetime ⟨*T* ⟩ in the population is proportional to 1*/*(*r* − *R*) (clearly, we must have *r > R*), indicating that populations with larger *R* have longer-lived individuals whose reproductive phase is limited to a smaller fraction of their lifespan. Note that while we allowed the reproduction rate to depend on age, here and in what follows we assumed that it does not depend on the state *χ* (for a discussion of the *P*_*R*_ = *P*_*R*_(*χ*) case see section IX in the SI).

The inheritance rule is defined by a Gaussian function that describes the probability that if a parent has a value *χ*^*′*^, its offspring will have a value *χ*. In our model this probability is a Gaussian of width *σ*, centered around *χ* = *χ*^*′*^:

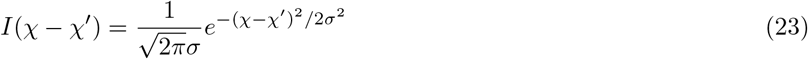

where *σ* measures the deviation from perfect inheritance. Such deviations from perfect inheritance can be due to epi-mutations that are encountered during the process of replication (“imperfect replication”).

By incorporating the modified *P*_*R*_(*T*) and *I*(*χ* −*χ*^*′*^) into the integral transform of eq. 5 and using the Taylor series expansion of the Fourier transform of *I*(*χ*−*χ*^*′*^) around small values of *kσ* (*k* and *χ* are conjugate Fourier transform variables - see section III in SI), we obtained steady state solutions for the population distribution and size in the case of a stationary environment. These solutions can be obtained by shifting the parameters *r* and *D* in eqs. 14 and 15: *r* → *r* − *R* and *D* → *D* + *rσ*^2^*/*2, respectively (SI section …). This yields:

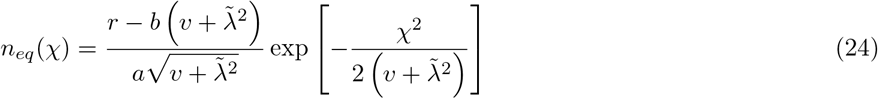

and

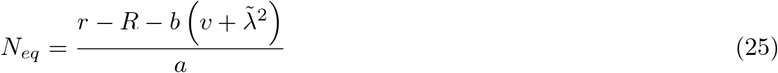

where

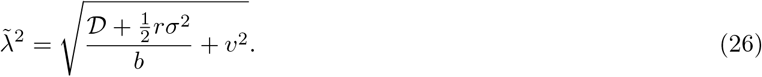

Since imperfect inheritance has the same effect as increasing the diffusion coefficient *D* by an amount that is proportional to *r*, the negative impact on population size cannot be compensated by increasing the rate of replication. Thus, unlike the case of perfect inheritance, the negative impact of imperfect inheritance on populations in a stationary environment can only be compensated by more effective reduction of the selective pressure within a generation (more negative *v*). The negative impact of age-dependent decline in fertility, on the other hand, can be readily compensated by higher rate of replication.

Additional insights into the effects of age-dependent decline in fertility and imperfect inheritance on populations that are exposed to changing environments can be obtained by expressing the differential equation for the distribution 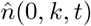 in terms of the following dimensionless parameters (SI, section VIII):

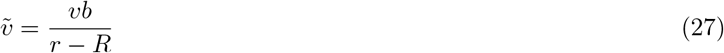

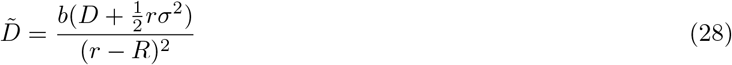

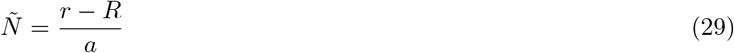

In terms of these dimensionless parameters, the equilibrium population in a stationary environment is given by

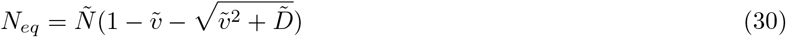

When measured in these dimensionless units, the time evolutions of different populations will be indistinguishable as long as they have the same 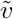 and 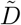 and the time is measured in units of *t*_*c*_ = 1*/*(*r* − *R*) (Fig. 5A and SI section VIII). The more complicated model with age-dependent decline in replication and imperfect inheritance can therefore be mapped to the simpler case of *R* = 0 and *σ* = 0, enabling extended analysis of how each of the various parameters (*r, R, v, D*, and *σ*) influence the time evolution of the population.

**FIG. 5:**
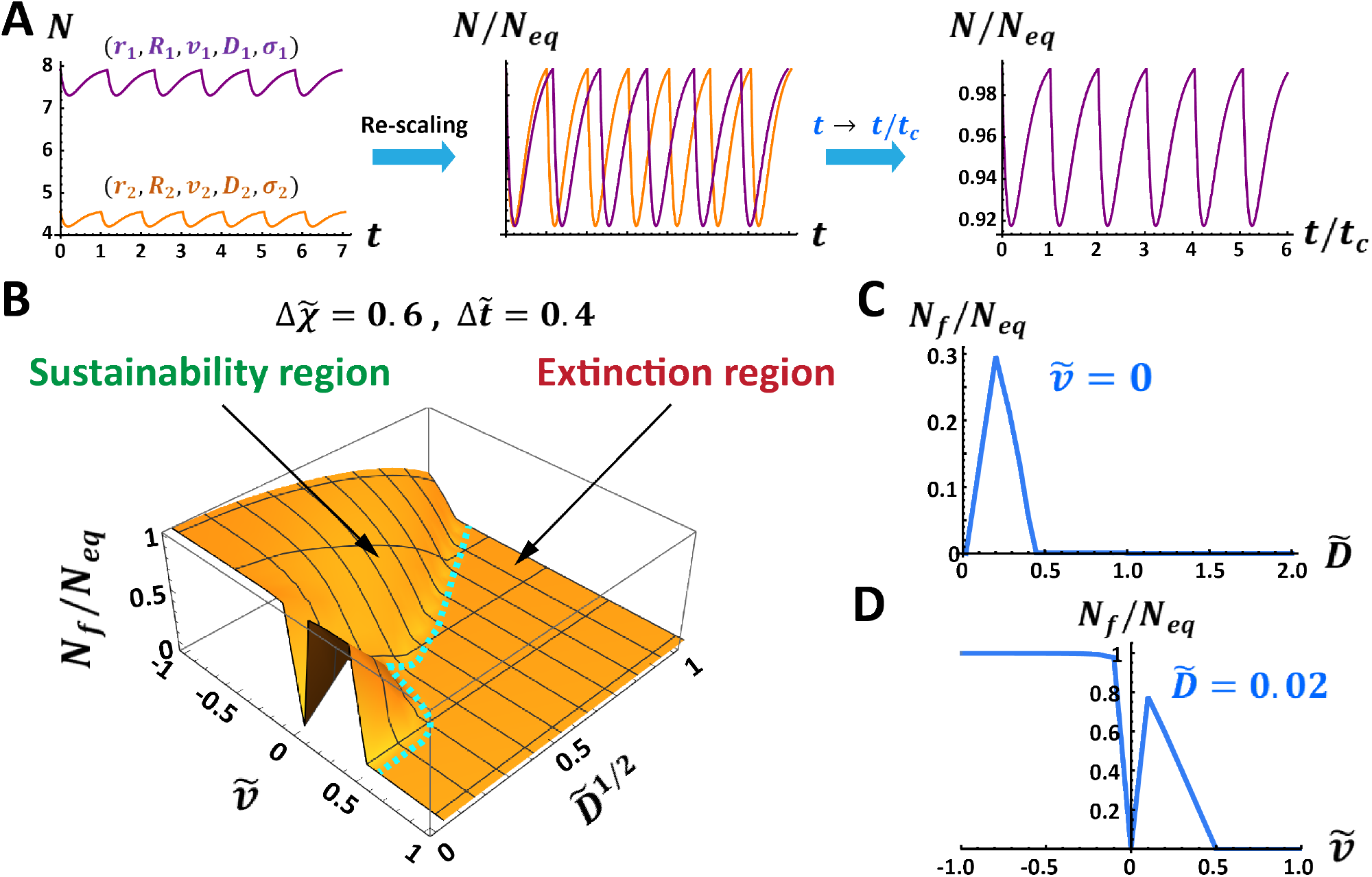
Long-term solutions to successive shifts of the environment (Type II dynamics), taking into consideration the effects of imperfect fidelity of inheritance and age-dependent decline in fertility. Comparison between populations under the same effective challenge, is enabled by normalizing the shift magnitude Δ*χ* and the duration between shifts Δ*t* by *χc* and *tc*. A) Effects of scaling the population size by *Neq* (middle) and measuring time in units of *t*_*c*_ (right). B) Landscape of *N*_*f*_ */N*_*eq*_ vs. 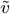 and 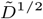, for fixed 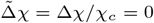 and 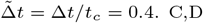. Sections of the landscape in (B) for 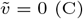 and 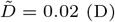, demonstrating significant impacts of changes that are within a generation on long-term population survival and size

Notwithstanding the above, it should be noted that for a given choice of Δ*t* and Δ*χ*, populations with broader distributions have a larger fraction of individuals experiencing a smaller increase in death rate (reduced increase in selective pressure). This means that the effective challenge acting on a population depends not only on the parameters of the environmental change (Δ*χ* and Δ*t*), but also on the equilibrium width of the distribution. This dependence can be eliminated by re-scaling Δ*χ* and Δ*t* by the characteristic width and time, 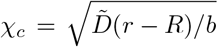 and *t*_*c*_ = 1*/*(*r*−*R*), respectively. Subsequent analysis of responses to environmental perturbation that is specified by the same 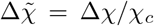 and 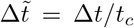 for all populations, enables investigation of how different choices of *r, v, D, R* and *σ* affect the steady state *N*_*f*_ */N*_*eq*_ under the same effective magnitude of challenge (defined by 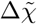 and 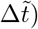. Fig. 5B displays the landscape of solutions *N*_*f*_ */N*_*eq*_ as a function of 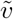 and 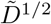 for the case of a unidirectional environmental change (Type II dynamics) specified by 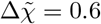 and 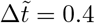 (see technical footnote [62]). It shows that the population-specific values of 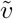 and 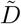 have a tremendous impact on the long-term evolution of the population. Note, in particular, that over a wide range of parameters, the sustainability of the population depends on having a sufficiently negative 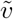 (Fig. 5B-D). The substantial impacts of 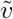 and 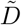 cannot be attributed to initial differences of the equilibrium distributions, because *N*_*f*_ is measured in units of the initial *N*_*eq*_ and the effect of population-specific differences in the width of the initial distribution has been eliminated by fixing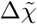 and 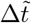.

Impacts of changing *r, R, v, D* and *σ* on *N*_*f*_ */N*_*eq*_ are deduced from Fig. 5B by determining the resulting values of the dimensionless parameters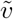 and 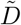 (Eqs. 28-30). Effects on the absolute size *N*_*f*_ (without normalization by *N*_*eq*_) can then be determined by taking into account the dependence of *N*_*eq*_ on 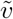 and 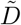 (Eq. 31).

To a first order approximation, a small increase in *R*, i.e. *R* → *R* + *δR* increases 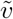 and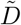 by *bvδR/*(*r*–*R*)^2^ and 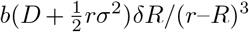, respectively, and decreases the equilibrium population *N*_*eq*_ by *δR/a*. Since a more negative *v* increases the positive impact of larger *R*, the benefit from reduction of selective pressure within a generation is more pronounced in populations of individuals exhibiting faster decline in fertility with age. This can be understood by realizing that these individuals reproduce more quickly, thus shortening the time to acquire beneficial changes that are transmitted to the offspring. Moreover, the landscape of *N*_*f*_ */N*_*eq*_ (Fig. 5B-D) shows that at the vicinity of 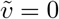 and 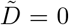, populations can also benefit from a small enough increase of 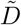 and even from sufficiently mild changes that increase the selective pressure within a generation (positive 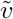). Since 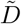 is also an increasing function of *D* and *σ*, similar benefits are conferred by a sufficiently small increase of either *D* or *σ* at the vicinity of 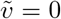 and 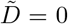. However, in contrast to increase in *D*, the negative impact of increasing *σ* cannot be compensated for by a larger rate of replication because the contribution of *σ* to 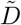 is scaled by *r*. This reveals a qualitative distinction between inheritance of stochastic changes that are acquired prior to proliferation (represented by *D*) and stochastic changes that occur during the replication process (represented by *σ*). As noted in the case of stationary environments, the negative impact of increased *σ* can nonetheless be compensated for by faster reduction of the selective pressure within a generation (more negative *v*).

One may also deduce the effects of changing parameters such as *R* while keeping the relative challenge fixed. For example, increasing *R* will lead to increases in 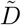 and in 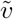. Accordingly, at different points in the diagram figure 5A, this could be either beneficial or harmful to the population.

## VIII. DISCUSSION

In contrast to genetic mutations that are relatively rare and can be transmitted over many generations, non-genetic changes of diverse kinds (such as epigenetic modifications) are frequently acquired within a lifetime but are generally far less stable than genetic variations. These changes include stochastic variations as well as non-random reactions to the environment that could potentially reduce or increase the rate of death (and/or reproduction) of individuals. Changes that reduce the rate of death in a new environment may benefit the population by rapidly decreasing the strength of selection; the impacts of inheriting these changes depend on the details and timescales of environmental changes. The long-term evolutionary implications of inheriting changes that are acquired within generation are further confounded by the limited persistence of these changes. Generating broadly applicable insights on long-term implications of inheriting non-genetic changes therefore requires theoretical modeling capable of drawing conclusions that are valid for most populations, despite their incredible diversity. Towards this goal, we formulated a population model that is independent of species-specific features and is based, instead, on rather general properties and capabilities whose impacts are investigated in the model, namely:

1. Age and state variables (*T* and *χ*) affecting the population dynamics.
2. Population parameters representing the magnitude of stochastic and directional changes in *χ* (*D* and *v*, respectively), the maximal rate of reproduction *r*, the rate of age-dependent decline in fertility *R*, and the deviation from perfect inheritance *σ* (also viewed as stochasticity of the reproductive process).
3. Functional dependence of the rates of death and proliferation *P*_*D*_ and *P*_*R*_, on the age and state variables, *T* and *χ* (the current study was limited to the cases in which *P*_*D*_ is independent of *T* and *P*_*R*_ is independent of *χ*).
4. Changes in *χ* that are proportional to the gradient of the death rate *P*_*D*_ (environmentally-driven reactions).
5. Inheritance of the state variable *χ* with variability that is defined by *σ*.
6. Environmental parameters representing carrying capacity *a*, and coefficient of selection *b* acting on the state variable, and the magnitude and rate of change of the adaptive peak of a type-dependent environment.

Previous models of population dynamics investigated population outcomes as a function of rates of reproduction, frequency of genetic mutations, population age structure, physiological state, and ecological factors in both stationary and dynamic environments [48–51, 53, 55, 63]. Since epigenetic inheritance of environmentally induced changes has been traditionally deemed infeasible, these models did not take this possibility into account. The accumulation of a large body of evidence supporting the feasibility of epigenetic inheritance prompted investigation of the adaptive value of non-genetic transmission [56]. Initial evaluation of the effect of transmitting spontaneous and environmentally induced states was based on a discrete model of a population with two states whose fitness is predefined for each of two environmental options [56, 57]. Since more realistic populations consist of individuals whose rates of death and birth depend on dynamic states and environments that are not fully specified in advance, the actual fitness is generally determined after the effect. The assessment of impacts of non-genetic transmission in such scenarios requires a modelling framework that can link arbitrary changes in the state and environment of individuals (at any moment) to longer-term impacts on the population. This was achieved by assigning every individual with a variable *χ*_*i*_(*t*) that can change within a lifetime and whose value at the time of proliferation is transmitted to the offspring (up to a small random deviation). In principle, *χ*_*i*_(*t*) can represent internal states (e.g. heritable changes in the epigenome and microbiome) as well as external states and structures that are created and/or modified by actions of individuals (e.g. niche construction). To consider state variables that are relevant for adaptation, we restricted the model to cases in which: (i) the death probability per unit time is a function of *χ*_*i*_, and (ii) the rate of directional change (drift) *f* (*x*_*i*_) is proportional to the gradient of this death term, 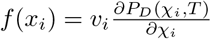.

In order to shift from individual- to population-level dynamics, we replaced the individual-specific drift and stochastic changes with a population average drift 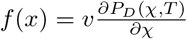 and diffusion *D*. This allowed us to derive an equation for the time evolution of the population distribution *n*(*χ, T, t*), subject to state-, age- and time-dependent rates of death *P*_*D*_(*χ, T, t*) and reproduction *P*_*R*_(*χ, T, t*) (in this work we focused on the case in which the reproduction rate does not depend on the state *χ* and the death rate does not depend on the age *T*). A drift that is proportional to the gradient of the death rate can represent qualitatively different scenarios of coping with the environment, corresponding respectively, to: (i) deterministic induction of a beneficial response that was previously selected because of its ability to reduce the rate of death, (ii) a non-beneficial response that is induced as a (deterministic) by-product of existing mechanisms and (iii) stochastic changes under constraints that reduce the likelihood of non-beneficial variations. In this case of constrained stochasticity, however, the drift does not apply at any moment, and is not even guaranteed. It should be thought of as a stochastic build-up of a directional change in *χ* under constraints that preferentially suppress changes in structures and mechanisms that are critical for survival. This biases the diffusion in *χ* towards the direction that reduces the death rate (equivalent to negative *v*). Since biased diffusion is qualitatively different from a deterministic drift, it cannot be formally modeled by the parameter *v*. It nonetheless gives rise to a ‘statistical drift’ that shares the benefit of moving away from deleterious changes. This alleviates the disadvantage of unbiased diffusion and may confer net benefit when the bias is large enough. While the overall benefit of this statistical drift is smaller than that of a deterministic drift with negative *v*, it has the unique advantage of enabling emergence of new adaptations within a lifetime [64]. As such, it extends the validity of the main insights of the model to the case of inherited adaptations that are newly forming during the individual’s lifetime (as opposed to adaptive responses that were selected in previous generations).

### Connecting short- and long-term effects of inheriting acquired changes

In the absence of migration of individuals, the overall population size can decrease only by the death of individuals and increase by birth events. While death, drift, and diffusion can occur at any age of individuals, the age of newborns at birth is always zero. This restriction is imposed by a boundary condition at *T* = 0, expressing the state distribution of newborns as an integral function of state- and age-dependent rate of the parental reproduction *P*_*R*_(*χ, T, t*) and the inheritance function *I*(*χ*−*χ*^*′*^). Given an initial distribution *n*(*χ, T, t*_0_), the time evolution of the population distribution is fully determined up to the time in which one of the population parameters (*r,R,D,v*, and *σ*) undergoes a significant change. Assuming that this time is similar to the timescale for acquiring genetic changes that drive noticeable effects at the population-level (typically over 10^3^ generations [65]), the model covers the population dynamics from an initial time point to many generations, thus enabling investigation of long-term effects of heritable states that are changing within a generation (1*/r*). In this work, we considered a simplified scenario in which the death rate has a quadratic dependence on the state variable *χ*, and the replication rate is either constant or age dependent. Impacts of directional and stochastic changes in state were represented, respectively, by drift that is proportional to the gradient of the death rate, 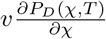 and diffusion of magnitude *D*. A drift with negative or positive *v* influences the rate of death by shifting the distribution towards or away from the adaptive peak, as well as by narrowing or broadening the distribution, respectively. The typical timescales for affecting the death rate by shifting and broadening the distribution are (*b*(|*v*|))^−1^ and 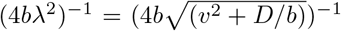, respectively. Parameters for which one of these timescales is shorter than 1*/r* enable investigation of long-term (multi-generational) effects of inheriting changes that are acquired within a generation. In this work, we focused on populations in either stationary environments, or two types of time-varying environments. Distinct impacts of the environment were represented by different components of the death rate, corresponding, respectively, to a global limitation of resources *aN* (*t*), and a force of selection *b*(*χ* −*χ*_0_(*t*))^2^ proportional to the quadratic distance from an adaptive peak *χ*_0_(*t*). The explicit time-dependence of *χ*_0_(*t*) defines temporal changes in the selective force due to temporal changes in the external environment, enabling investigation of impacts of epigenetic inheritance under different patterns and timescales of environmental challenge.

### Impacts of epigenetic inheritance in stationary and dynamic environments

The benefits or disadvan-tages of the drift depend on the sign and magnitude of *v*, as well as the type and dynamics of the external environment. Under stationary environments and non-vanishing variability (*D >* 0), the steady state population size increases as a function of negative *v* and decreases with positive *v*. These impacts are caused, respectively, by ongoing reduction or increase in the rate of death without affecting the reproduction rate. Negative *v* also reduces the time for population relaxation after a single (or a sufficiently rare) transition to another stationary environment. This is mediated by the directional shift of the distribution towards the new adaptive peak. A shift in the opposite direction (positive *v*), however, does not always increase the relaxation time. When *v* and *D* are sufficiently small, an increase in *v* has the counter-intuitive effect of shortening the time for relaxation. This effect is caused by contribution of positive *v* to broadening of the distribution, which in turn increases the fraction of individuals whose state *χ* is closer to the new adaptive peak (as long as the width of distribution is below a certain threshold). For small enough *v*, this effect exceeds the negative impact of shifting the distribution away from the adaptive peak. Since the distribution is also broadened by diffusion, the relaxation time in this regime is also shortened by an increase in *D*. Beyond this regime, however, larger values of either *D* or (positive) *v* increase the time for relaxation by further widening the distribution and/or shifting it away from the adaptive peak.

Under periodically switching, as well as unidirectionally shifting environments, we found that negative *v* generally contributes to population recovery, and may even be essential to avoid extinction in the presence of a strong challenge. In a scenario of unidirectional shifts, the combination of negative *v* and transmission of the acquired state confers two obvious benefits: reduction of death rate within a generation, and persistence of the lower death rate in newborns. In the case of periodic switching, however, the effect of the transmission itself depends on the time of environmental switching and is not necessarily beneficial. If the switching time is larger than a few generations, the transmission is mostly beneficial (as noted by the earlier work of Lachmann and Jablonka [57]). On the other hand, when Δ*t* is comparable to the generation time, newborns often inherit *χ* that is farther away from the new adaptive peak, thus contributing to an initially elevated rate of death. Nonetheless, when *v* is negative, the deleterious impact of transmitting a maladaptive *χ* is counteracted by subsequent shift of the distribution towards the new adaptive peak (as well as by narrowing the distribution around the peak). Analysis of the long-term size of the recovered population showed that the net effect is positive even when the switching time of the environment equals the generation time 1*/r* (Fig. 3D). A sufficiently negative *v* also enables the sustainability of the population in a range of Δ*χ* and *D* that would otherwise result in extinction. Escape from extinction may also be achieved by faster reproduction (larger *r*) of populations with large 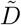 and 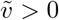. This is indicated by the decrease of 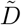 and 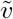 as a function of *r* (Eqs. 28-29). In any real organism, however, the rate of reproduction cannot assume arbitrarily large levels without compromising survival (increasing *P*_*D*_) and lowering the accuracy of reproduction (increasing *σ*). Accordingly, the ability to avoid extinction by faster reproduction is limited to a certain range of *r*. This contrasts with the largely unrestricted benefits of reducing the selective pressure within a lifetime (more negative *v*), which include: larger population sizes under stationary environments (Eq. 30), expedited recovery from a sudden transition to a challenging environment (Fig. 2), avoiding extinction within a range of highly challenging scenarios (indicated by the survival condition of Eq. 20), and increased size of the recovered population relative to the equilibrium size in a stationary environment (demonstrated by the landscape of *N*_*f*_ */N*_*eq*_, either as a function of *v* and *D*^1*/*2^ in Figs. 3,4, or 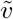 and 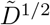 in Fig. 5).

Traditional consideration of heritable stochastic variations was restricted to the case of rare mutations, typically represented by a constant rate of mutagenesis with no distinction of when the mutations are generated. Extending this to stochastic variations that are frequently encountered within a lifetime and inherited with imperfect fidelity, reveals a significant distinction between the influence of stochastic variations that are encountered prior to reproduction (represented by *D*) versus that of stochastic transmission in the process of reproduction (represented by *σ*). While both are generally deleterious (except for a narrow range of parameters), the negative impact of variations that are acquired prior to reproduction is independent of the replication rate, whereas the negative impact of variations in transmission scales with the reproduction rate. This gives rise to unique implications of imperfect inheritance, namely: (i) A penalty on reproduction (manifested as an effective increase in *D* by 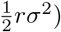, and (ii) limited ability to compensate for the deleterious impact of imperfect inheritance by increasing the rate of reproduction. The resulting limitations on the benefit gained by faster reproduction provide an advantage for populations exhibiting higher fidelity of epigenetic inheritance. The manifestation of this advantage in real populations is limited by the mechanistic difficulty of transmitting acquired changes. This difficulty depends on the complexity of the organism and the type of inherited change. With that in mind, it is interesting to notice that the fidelity of inheritance is typically high when the barrier for transmission is low. Obvious examples include inheritance of cytoplasmic factors by cell division in unicellular organisms, inheritance of the gut microbiome composition in animals and cultural inheritance in humans.

The findings of this study were restricted to a baseline model comprising: a single state variable *χ* that is influenced by the gradient of the death rate, an exponentially decreasing rate of reproduction with age *T*, and a rate of death that depends on both *χ* and *t*. The baseline formulation was chosen for simplicity (e.g., existence of an analytical solution of the model) and its main conclusions were found to hold for a more realistic age-dependence of the reproduction rate (an increase from zero at birth to a maximum at an intermediate age, followed by exponential decay at old age; SI, section X). The underlying modelling framework (equations 4-7) is readily extendable to rates of death and reproduction that are arbitrary functions of age, state, and time (example with state-dependent rate of reproduction is provided in SI, section IX). However, these extensions are typically not amenable to analytic solutions, thus requiring numerical analysis. While the solutions in these cases depend on the specific formulation of the death and reproduction rates, the inheritance of changes that are acquired within a generation is expected to have long-term effects in a wide range of formulations.

## Supporting information

Additional Details for Derivations

## ACKNOWLEDGMENTS

This work was supported by the Sir John Templeton Foundation (grant numbers: 40663 and 61122). Helpful discussions with D. Kessler and N. Schnerb are gratefully acknowledged.

## References

[1] A. Bosković and O. Rando, Transgenerational epigenetic inheritance, Annu. Rev. Genet. 52, 21–41 (2018).

[2] G. Cavalli and E. Heard, Advances in epigenetics link genetics to the environment and disease, Nature 571, 489 (2019).

[3] M. Fitz-James and G. Cavalli, Molecular mechanisms of transgenerational epigenetic inheritance, Nat Rev Genet. 23, 325 (2022).

[4] M. Miryeganeh and H. Saze, Epigenetic inheritance and plant evolution, Population Ecology 62, 17 (2020).

[5] I. Lacal and R. Ventura, Epigenetic inheritance: Concepts, mechanisms and perspectives, Front Mol Neurosci. 11, 292 (2018).

[6] G. Cavalli and R. Paro, The Drosophila Fab-7 chromosomal element conveys epigenetic inheritance during mitosis and meiosis, Cell 93, 505–518 (1998).

[7] H. Morgan, H. Sutherland, D. Martin, and E. Whitelaw, Epigenetic inheritance at the agouti locus in the mouse, Nature Genetics 23, 314–318 (1999).

[8] M. Anway, A. Cupp, M. Uzumcu, and M. Skinner, Epigenetic transgenerational actions of endocrine disruptors and male fertility, Science 308, 1466 (2005).

[9] O. Rechavi, G. Minevich, and O. Hobert, Transgenerational inheritance of an acquired small RNA-based antiviral response in C. elegans, Cell 146 (1248–1256).

[10] A. Klosin, E. Casas, C. Hidalgo-Carcedo, T. Vavouri, and B. Lehner, Transgenerational transmission of environmental information in C. elegans, Science 356, 320 (2017).

[11] S. Ng, R. Lin, D. Laybutt, R. Barres, J. Owens, and M. Morris, Chronic high-fat diet in fathers programs beta-cell dysfunction in female rat offspring, Nature 467, 963–966 (2010).

[12] B. Carone, L. Fauquier, N. Habib, J. Shea, C. Hart, R. Li, C. Bock, C. Li, H. Gu, P. Zamore, A. Meissner, Z. Weng, H. Hofmann, N. Friedman, and O. Rando, Paternally induced transgenerational environmental reprogramming of metabolic gene expression in mammals, Cell 143, 1084 (2010).

[13] O. Rechavi, L. Houri-Ze’evi, S. Anava, W. Goh, S. Kerk, G. Hannon, and O. Hobert, Starvation-induced transgenerational inheritance of small RNAs in C. elegans, Cell 158, 277 (2014).

[14] D. Beck, M. Maamar, and M. Skinner, Integration of sperm ncRNA-directed DNA methylation and DNA methylation-directed histone retention in epigenetic transgenerational inheritance, Epigenetics and Chromatin 14, 6 (2021).

[15] S. King and M. Skinner, Epigenetic transgenerational inheritance of obesity susceptibility, Trends Endocrinol Metab. 31, 478 (2020).

[16] B. Dias and K. Ressler, Parental olfactory experience influences behavior and neural structure in subsequent generations, Nat Neurosci. 17, 89 (2014).

[17] J. Kelley, M. Tobler, D. Beck, I. Sadler-Riggleman, C. Quackenbush, L. Arias Rodriguez, and M. Skinner, Epigenetic inheritance of dna methylation changes in fish living in hydrogen sulfide-rich springs, Natl Acad Sci U S A 26 (2021).

[18] L. Goldberg and T. Gould, Multigenerational and transgenerational effects of paternal exposure to drugs of abuse on behavioral and neural function, Eur J Neurosci. 50, 2453 (2019).

[19] C. Guerrero-Bosagna, M. Settles, B. Lucker, and M. Skinner, Epigenetic transgenerational actions of vinclozolin on promoter regions of the sperm epigenome, PLoS One 5, 13100 (2010).

[20] A. Holloway, D. Cuu, K. Morrison, H. Gerstein, and M. Tarnopolsky, Transgenerational effects of fetal and neonatal exposure to nicotine, Endocrine 31, 254 (2007).

[21] J. Švorcová, Transgenerational epigenetic inheritance of traumatic experience in mammals, Genes 14, 120 (2023).

[22] F. Bantignies, C. Grimaud, S. Lavrov, and G. Cavalli, Inheritance of Polycomb-dependent chromosomal interactions in Drosophila, Genes and Development 17, 2406 (2003).

[23] K. Seong, D. Li, H. Shimizu, R. Nakamura, and S. Ishii, Inheritance of stress-induced, ATF-2-dependent epigenetic change, Cell 145, 1049 (2011).

[24] A. Ashe, S. A, W. EM, M. J, B. MP, C. AC, D. AL, L. Goldstein, N. Lehrbach, J. Le Pen, G. Pintacuda, A. Sakaguchi, P. Sarkies, S. Ahmed, and E. Miska, piRNAs can trigger a multigenerational epigenetic memory in the germline of C. elegans, Cell 150, 88 (2012).

[25] A. Weyrich, S. Yasar, D. Lenz, and J. Fickel, Tissue-specific epigenetic inheritance after paternal heat exposure in male wild guinea pigs, Mamm Genome 31, 157 (2020).

[26] S. Wang, K. Lau, K. Lai, J. Zhang, A. Tse, J. Li, Y. Tong, T. Chan, C. Wong, J. Chiu, D. Au, A. Wong, R. Kong, and R. Wu, Hypoxia causes transgenerational impairments in reproduction of fish, Nat Commun. 7, 12114 (2016).

[27] S. Stern, O. Snir, E. Mizrachi, M. Galili, I. Zaltsman, and Y. Soen, Reduction in maternal Polycomb levels contributes to transgenerational inheritance of a response to toxic stress in flies, J Physiol. 592, 2343 (2014).

[28] Y. Fridmann-Sirkis, S. Stern, M. Elgart, M. Galili, A. Zeisel, N. Shental, and Y. Soen, Delayed development induced by toxicity to the host can be inherited by a bacterial-dependent, transgenerational effect, Front Genet. 5, 27 (2014).

[29] M. Elgart, S. Stern, O. Salton, Y. Gnainsky, Y. Heifetz, and Y. Soen, Impact of gut microbiota on the fly’s germ line, Nat Commun. 15, 11280 (2016).

[30] R. Bonduriansky, A. Crean, and T. Day, The implications of nongenetic inheritance for evolution in changing environments, Evolutionary Applications 5, 192–201 (2011).

[31] T. Pentinat, M. Ramon-Krauel, J. Cebria, R. Diaz, and J. Jimenez-Chillaron, Transgenerational inheritance of glucose intolerance in a mouse model of neonatal overnutrition, Endocrinology 151, 5617 (2010).

[32] P. Sarkies, Molecular mechanisms of epigenetic inheritance: Possible evolutionary implications, Semin Cell Dev Biol. 97, 106 (2020).

[33] A. Ashe, V. Colot, and B. Oldroyd, How does epigenetics influence the course of evolution?, Philos Trans R Soc Lond B Biol Sci. 376, 20200111 (2021).

[34] C. Richards, K. Verhoeven, and O. Bossdorf, <u>Evolutionary Significance of Epigenetic Variation</u>, Plant Genome Diversity, Vol. 1 (Springer, 2012) p. 257–274.

[35] E. Jablonka, The evolutionary implications of epigenetic inheritance, Interface Focus. 7, 20160135 (2017).

[36] M. Schlaepfer, M. C. Runge, and P. W. Sherman, Ecological and evolutionary traps, Trends in Ecology and Evolution 17, 474 (2002).

[37] W. Burggren, Dynamics of epigenetic phenomena: intergenerational and intragenerational phenotype ‘washout’, J Exp Biol. 218, 80 (2015).

[38] T. Burton, I. Ratikainen, and S. Einum, Environmental change and the rate of phenotypic plasticity, Glob Chang Biol. 28, 5337 (2022).

[39] D. Osmanovic, D. Kessler, Y. Rabin, and Y. Soen, Darwinian selection of host and bacteria supports emergence of Lamarckian-like adaptation of the system as a whole, Biol Direct. 13, 24 (2018).

[40] J. Roughgarden, Holobiont evolution: Population theory for the hologenome, Am Nat. 201, 763 (2023).

[41] S. Hormoz, N. Desprat, and B. Shraiman, Inferring epigenetic dynamics from kin correlations, Proc Natl Acad Sci U S A 112, E2281 (2015).

[42] M. Lind and F. Spagopoulou, Evolutionary consequences of epigenetic inheritance, Heredity 121, 205 (2018).

[43] R. Fox, J. Donelson, C. Schunter, T. Ravasi, and J. Gaitaán-Espitia, Beyond buying time: the role of plasticity in phenotypic adaptation to rapid environmental change, Phil. Trans. R. Soc. B 374, 20180174 (2019).

[44] I. Overcast, M. Ruffley, J. Rosindell, L. Harmon, P. Borges, B. Emerson, R. Etienne, R. Gillespie, H. Krehenwinkel, D. Mahler, F. Massol, C. Parent, J. Patinño, B. Peter, B. Week, C. Wagner, M. Hickerson, and A. Rominger, A unified model of species abundance, genetic diversity, and functional diversity reveals the mechanisms structuring ecological communities, Mol Ecol Resour. 21, 2782 (2021).

[45] O. Savolainen, M. Lascoux, and J. Merilä, Ecological genomics of local adaptation, Nat Rev Genet. 14, 807 (2013).

[46] W. Lowe, R. Kovach, and F. Allendorf, Population genetics and demography unite ecology and evolution, Trends in Ecology and Evolution 32, 141 (2017).

[47] G. Rehfeldt, L. Leites, D. Joyce, and A. Weiskittel, Role of population genetics in guiding ecological responses to climate, Glob Chang Biol. 24, 858 (2018).

[48] E. Kussell and S. Leibler, Phenotypic diversity, population growth and information in fluctuating environments, Science 309, 2075 (2005).

[49] A. de Roos, A gentle introduction to physiologically structured population models, in <u>Structured-Population Models in Marine, Terrestrial, and Freshwater Systems</u>, edited by S. Tuljapurkar and H. Caswell (Springer US, Boston, MA, 1997) p. 119—204.

[50] H. Inaba, <u>Age-structured population dynamics in demography and epidemiology</u> (Springer, 2017).

[51] L. Tsimring and H. Levine, RNA virus evolution via a fitness-space model, Phys. Rev. Lett. 76, 4440 (1996).

[52] N. Schnerb and D. Nelson, Non-Hermitian localization and population biology, Phys. Rev. E 58, 1383 (1998).

[53] D. Kessler and H. Levine, Phenomenological approach to cancer cell persistence, Phys. Rev. Lett. 129, 108101 (2022).

[54] M. J. Melissa, B. H. Good, D. S. Fisher, and M. M. Desai, Population genetics of polymorphism and divergence in rapidly evolving populations, Genetics 221, 1 (2023).

[55] D. Kessler and N. Shnerb, Neutral-like abundance distributions in the presence of selection in a continuous fitness landscape, Journal of Theoretical Biology 345, 1 (2014).

[56] E. Jablonka, The adaptive advantage of phenotypic memory in changing environments, Phil. Trans. R. Soc. Lond. B 350, 133 (1995).

[57] M. Lachmann and E. Jablonka, The inheritance of phenotypes: an adaptation to fluctuating environments, J. Theor. Biol. 181, 1 (1996).

[58] B. Keyfitz and N. Keyfitz, The McKendrick partial differential equation and its uses in epidemiology and population study, Mathematical and Computer Modelling 26, 1 (1997).

[59] A. McKendrick, Applications of mathematics to medical problems, Proc. Edinb. Math. Soc. 44, 98 (1926).

[60] H. von Forester, Some remarks on changing populations, in <u>The Kinetics of Cellular Proliferation</u>, edited by F. Stahlman (Grune and Stration New York and London, 1959) pp. 382–407.

[61] L. Landau and E. Lifshitz, <u>Statistical Physics, Part 1</u> (Butterworth-Heinemann, Oxford, 1980).

[62] The population is represented in terms of Neq rather than Ñ. As Neq is a linear function of Ñ, Neq can be substituted when measuring population scales.

[63] I. Akushevich, A. Yashkin, J. Kravchenko, F. Fang, K. Arbeev, F. Sloan, and A. Yashin, A forecasting model of disease prevalence based on the McKendrick-von Foerster equation, Math Biosci. 311, 31 (2019).

[64] Y. Soen, M. Knafo, and M. Elgart, A principle of organization which facilitates broad Lamarckian-like adaptations by improvisation, Biol. Direct 10, 68 (2015).

[65] O. Rando and K. Verstrepen, Cell 128, 655 (2007).

