## Additional Details for Derivations for "Population model of epigenetic inheritance of acquired adaptation to changing environments"

Dino Osmanović*

*Department of Mechanical and Aerospace Engineering,*

*University of California, Los Angeles, Los Angeles 90095, CA, USA*

Yitzhak Rabin

*Department of Physics, Bar-Ilan University, Ramat Gan 5290002, Israel*

Yoav Soen

*Department of Bio-molecular Sciences,*

*Weizmann Institute of Science, Rehovot, Israel*

(Dated: February 15, 2024)

### CONTENTS

I. Derivation of the extended McKendrick–von Foerster equation 2

II. Symbiosis leading to effective drift 4

III. Solution for $R = 0, I = \delta(\chi - \chi')$ model 6

IV. Stationary Distribution 9

V. Adaptation to a Sudden change in the environment 10

VI. Continual variation of the environment 12

VII. Conditions on survival 14

VIII. Fertility, Inheritance and their effect on adaptation to environment 15

IX. $\chi$-dependent replication 18

*

X. More realistic birth rates 19

References 21

### I. DERIVATION OF THE EXTENDED MCKENDRICK–VON FOERSTER EQUATION

We shall extend the McKendrick–von Foerster equation to include the effect of changes, both pre-programmed and random, that take place during the lifetime of individuals and affect their survival probabilities (epigenetics, microbiome, etc.)

The equations that govern the time ($t$) evolution of the distribution of ages $T$ in a population $n(T,t)$ are given by:

$$\frac{\partial n\left(T,t\right)}{\partial t} = -\frac{\partial n\left(T,t\right)}{\partial T} - P_D\left(T,t\right) n\left(T,t\right) \tag{1}$$

$$n\left(0,t\right) = \int P_R\left(T',t\right) n\left(T',t\right) \mathrm{d}T' \tag{2}$$

$$n\left(\infty,t\right) = 0, \tag{3}$$

which can be derived by considering the fundamental processes involved. Here $P_D\left(T,t\right)$ and $P_R\left(T',t\right)$ are the death and replication rates, respectively. Note that death is incorporated as a sink term in Eq. 1 and replication enters as an initial condition (at age $T=0$), in Eq. 2.

We shall now derive the extended equation by including an additional degree of freedom in this equation. To begin, we consider an extended distribution, which includes both age $T$ and some other parameter $\chi$.

$$n\left(T,\chi,t\right) = \sum_{i=1}^{N} \delta\left(T - T_i(t)\right) \delta\left(\chi - \chi_i(t)\right) \tag{4}$$

where after a period of time $\Delta t$, we have:

$$n\left(T,\chi,t+\Delta t\right) = \sum_{i=1}^{N} \delta\left(T - T_i(t+\Delta t)\right) \delta\left(\chi - \chi_i(t+\Delta t)\right) \tag{5}$$

In order to convert this into a differential equation, we need to specify update rules for the parameters $T$ and $\chi$. For each individual $i$, the update rule for age in time is simply:

$$T_i(t+\Delta t) = T_i(t) + \Delta t \tag{6}$$

For the update of the additional parameter $x_i$ we choose a drift-diffusion scheme:

$$\chi_i(t+\Delta t) = \chi_i(t) + f(\chi_i, t)\Delta t + \xi_i(t) \tag{7}$$

where $\xi_i$ is a random number of mean 0 with correlation function $\langle \xi_i(t)\xi_j(t')\rangle = 2D\delta_{i,j}\delta(t-t')$, and $f(\chi_i, t)$ is a deterministic change in $\chi_i$ per unit time (here and in the following we assume that $f$ does not depend on the age of the individual, $T_i$). Note that the parameter $\chi_i$ changes *during the lifetime* of the individual. In the following we assume that both the form of the deterministic function $f$ and the statistical properties (e.g., variance) of the random variable $\xi$, are the same for all individuals in the population.

When combined, these update rules correspond to the following Fokker-Planck like equation that we will refer to as the extended McKendrick–von Foerster equation:

$$\frac{\partial n\left(T,\chi,t\right)}{\partial t} = -\frac{\partial n\left(T,\chi,t\right)}{\partial T} - \frac{\partial}{\partial \chi}\left(f(\chi)n(T,\chi,t)\right) + D\frac{\partial^2 n(T,\chi,t)}{\partial \chi^2} - P_D\left(T,\chi,t\right)n\left(T,\chi,t\right) \tag{8}$$

This equation corresponds to individuals aging in time, with some other parameter $\chi$ of the individuals evolving according to the drift-diffusion equation, with a drift term $f(\chi, t)$ that depends itself on the parameter $\chi$. This state parameter could correspond to any facet of the individual. Note that the death rate $P_D(T, \chi, t)$ is, in general, a function of both the age and state parameters $T$ and $\chi$, respectively, and of time $t$.

The replication rule is more complicated. It is given by:

$$n(0,\chi,t) = \int P_R(T',\chi,t) I(\chi-\chi') n(T',\chi',t) \mathrm{d}\chi' \mathrm{d}T' \tag{9}$$

This equation is similar to Eq. 1, except now we have an additional term accounting for the inheritance of the state parameter $\chi$, given by the function $I(\chi - \chi')$. For example, if the inheritance function is given by $I(\chi - \chi') = \delta(\chi - \chi')$ this corresponds to exact inheritance of the parameter $\chi$, which changes during the parent's lifetime. The possibility of imperfect inheritance can be modelled by a Gaussian inheritance function.

We therefore arrive at the modified McKendrick–von Foerster equation (with its boundary conditions at $T = 0$ and $T = \infty$):

$$\frac{\partial n\left(T,\chi,t\right)}{\partial t} = -\frac{\partial n\left(T,\chi,t\right)}{\partial T} - \frac{\partial}{\partial \chi}\left(f(\chi)n(T,\chi,t)\right) + D\frac{\partial^2 n(T,\chi,t)}{\partial \chi^2} - P_D(T,\chi)n(T,\chi,t) \tag{10}$$

$$n(0,\chi,t) = \int P_R(T',\chi',t)I(\chi-\chi')n(T',\chi',t)\mathrm{d}\chi'\mathrm{d}T' \tag{11}$$

$$n(\infty,\chi,t) = 0 \tag{12}$$

### II. SYMBIOSIS LEADING TO EFFECTIVE DRIFT

We shall in the following motivate particular choices of the drift function $f(\chi)$ appearing in the main text by considering a system of two populations co-evolving under the following assumptions:

- One population breeds much faster than the other. We call the fast breeders "bacteria" and the slow breeders "hosts"
- A population of the bacteria is associated with an individual host. In a population of hosts, we have a population of populations of bacteria, each one corresponding to a particular host, i.e. a microbiome.
- The probability of survival of each host depends, in some sense, on the properties of its bacterial population. For example, if we take some set of genomic properties of the fast population, the survival (or death) probability of the slow breeder will depend on these properties.

One particularly attractive example would correspond to a system where the death probability of the host depends on the mean $\langle\chi\rangle$ of some set of bacterial properties. We can use equation 10 with $f(\chi) = 0$ to construct a Darwinian model of the bacteria under some environment (defined through some parameters that characterize the dependence of the death probability $P_D$ on $\chi$).

If the mean of the bacterial population $\langle\chi\rangle$ is away from the minimum of the death rate $P_D(\chi) \sim \chi^2$, the mean will evolve in time until it reaches the minimum. When the dynamics of the mean are observed in this simple scenario, they are seen to be exponential (see fig 1B). The importance of this observation is that if we look at the process on the timescale of the host, it would appear as if it is responding to its environment by changing its value of $\langle\chi\rangle$ by introducing a drift function of the following form:

$$f(\chi) = v\frac{\partial P_D(\chi)}{\partial \chi} \tag{13}$$

In other words, an organism with parameter $\langle\chi\rangle$ responds to environment, by moving away from regions of large death rate for negative $v$ and by moving towards these regions for positive $v$. We stress that this arises only by considering coupled co-evolution over two divergent time scales. When $v < 0$, the evolutionary objectives of both host and bacteria are aligned and the bacterial adaptation proceeds in a direction which lowers the death rate of the host. However, the opposite can be the case where $v > 0$. In this scenario, the direction of bacterial adaptation leads to greater death rates of the host, as would be the case for pathogenic bacteria.

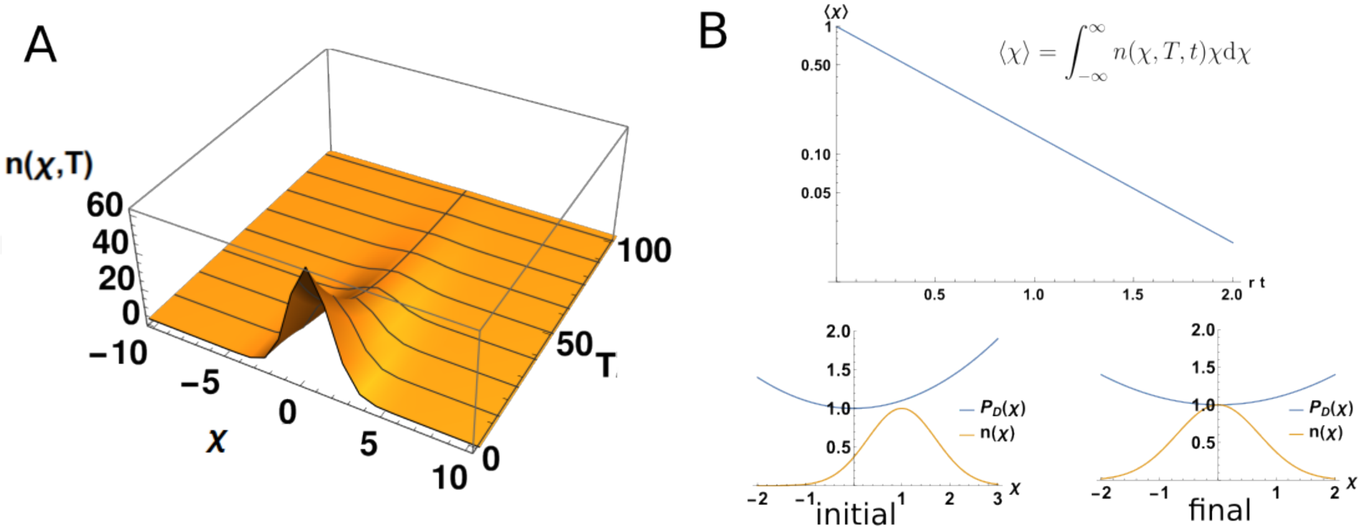


FIG. 1. Fundamental elements of the mathematical model. We construct expressions for how the distribution $n(\chi, T, t)$ changes in time. In panel A) we show an example distribution for an environment in which the death rate scales as $\sim \chi^2$, with $D = 1, v = 0, r = 2, a = 1, b = 1$. This leads to a steady state population distribution that is exponentially declining with age $T$ and is Gaussian in the state parameter $\chi$. B) The change in time of the mean of an initially misaligned initial population, subject to only diffusive dynamics. The mean relaxes exponentially, which can be interpreted as drift, on a longer timescale. (a=1,b=1,D=1,v=0,r=1)

### III. SOLUTION FOR $R = 0, I = \delta(\chi - \chi')$ MODEL

We will in the following focus on an analytical example; this requires introducing Fourier transforms in $\chi$ and Laplace transforms in $T$. We will use the form of the death rate that we have introduced in the main text:

$$P_D(\chi) = a \int \mathrm{d}\chi' \mathrm{d}T' n(T', \chi', t) + b\chi^2 \tag{14}$$

This form of death rate has a term proportional to the total amount of individuals (global feedback due to population) and a term that depends quadratically on $\chi$ and ensures that the death rate has a minimum at $\chi = 0$ ("harmonic landscape"). Here and in the following we assume that the death rate does not depend on the age parameter and is a function of the state parameter $\chi$ only. Notice that while this simplyfying assumption appears to be unphysical, even in the case of a constant death rate the population decreases exponentially with age. Therefore, the expected exponential increase of the death rate with $T$ for ages exceeding some value $T*$ would act as a cutoff of the population distribution at $T*$ and will have a negligible effect on our results.

The additional assumption that was mentioned in the main text is that of perfect replication, $I(\chi - \chi') = \delta(\chi - \chi')$. This corresponds to offspring having exactly the same value of $\chi$ as their parents. The replication rate is given by:

$$P_R(T) = r \exp(-RT) \tag{15}$$

where $r$ is the maximal replication rate and $1/R$ defines a time scale on which the ability to replicate decreases with age. Here and in the following we assume that the replication rate does not depend on the state parameter $\chi$ and is time-independent (i.e., does not depend on the changing environment).

As our starting point, we apply a Laplace transform in age $T$, and a Fourier transform in variable $\chi$ to the population distribution $n(T, \chi, t)$:

$$\hat{n}(S, k, t) = \int_{-\infty}^{\infty} \mathrm{d}\chi \exp(-ik\chi) \int_{0}^{\infty} \mathrm{d}T \exp(-ST) n(T, \chi, t) \tag{16}$$

We note here that the total number of individuals is given by $\hat{n}(0, 0, t)$. Applying the transforms to Eq. 10 yields:

$$\frac{\partial\hat{n}(S,k,t)}{\partial t} = -S\hat{n}(S,k,t) + r\hat{n}(R,k,t) + 2vbk\frac{\partial\hat{n}(S,k,t)}{\partial k} - Dk^2\hat{n}(S,k,t) - a\hat{n}(S,k,t)\hat{n}(0,0,t) + b\frac{\partial^2\hat{n}(S,k,t)}{\partial k^2} \tag{17}$$

For now, we ignore the fact that the total population $\hat{n}(0,0,t)$ is a functional of $\hat{n}(S,k,t)$), and treat it as some given function of time (we shall later demand that the solution is self-consistent in order to reintroduce this feedback). Furthermore, we ignore the age dependence of the replication time of the individual, by setting $R = 0$. In this case the entire behavior of the system is driven by the $S = 0$ mode in Laplace space, that obeys the following equation:

$$\frac{\partial\hat{n}(0,k,t)}{\partial t} = r\hat{n}(0,k,t) + 2vbk\frac{\partial\hat{n}(0,k,t)}{\partial k} - Dk^2\hat{n}(0,k,t) - a\hat{n}(0,k,t)N(t) + b\frac{\partial^2\hat{n}(0,k,t)}{\partial k^2} \tag{18}$$

Let us separate $\hat{n}(0,k,t)$ as $K(k)L(t)$, this yields (after dividing through by $\hat{n}$):

$$\frac{1}{L(t)}\frac{\partial L(t)}{\partial t} + aN(t) - r = +2vbk\frac{1}{K(k)}\frac{\partial K(k)}{\partial k} - Dk^2 + b\frac{1}{K(k)}\frac{\partial^2 K(k)}{\partial k^2} \tag{19}$$

Having successfully separated the variables here, we can solve each side being equal to some constant $-\mu$. This yields:

$$L(t) = A\exp\left(\int_0^t \mathrm{d}t'(r - aN(t') - \mu)\right) \tag{20}$$

$$K(k) = Be^{-\frac{1}{2}k^2\sqrt{\frac{bv^2+\mathcal{D}}{b}} - \frac{k^2v}{2}}H_{\frac{-vb-\sqrt{\frac{bv^2+\mathcal{D}}{b}}b+\mu}{2b\sqrt{\frac{bv^2+\mathcal{D}}{b}}}}\left(k\sqrt[4]{\frac{bv^2+\mathcal{D}}{b}}\right) \tag{21}$$

where $H_m(x)$ is a Hermite polynomial of rank $m$ and $A$ and $B$ are some constants. As these polynomials form of complete set, we set the degree of the polynomial equal to an integer $m$ and solve for $\mu$, giving $\mu = b\left((2m+1)\sqrt{\frac{\mathcal{D}}{b} + v^2} + v\right)$, and we can write down the complete solution:

$$\hat{n}(0,k,t) = \sum_m c_m e^{\left(\int_0^t \mathrm{d}t'\left(r - aN(t') - b\left((2m+1)\sqrt{\frac{\mathcal{D}}{b}+v^2}+v\right)\right)\right)} e^{-\frac{1}{2}k^2\left(\sqrt{\frac{\mathcal{D}}{b}+v^2}+v\right)} H_m\left(k\sqrt[4]{v^2 + \frac{\mathcal{D}}{b}}\right) \tag{22}$$

We see a common factor appearing, $\sqrt{\frac{\mathcal{D}}{b} + v^2}$, which we define as

$$\lambda^2 = \sqrt{\frac{\mathcal{D}}{b} + v^2} \tag{23}$$

for brevity.

Using the orthogonality of the Hermite polynomials allows us to write down the constants $c_m$:

$$c_m = \frac{\lambda}{\sqrt{\pi} 2^m m!} \int \mathrm{d}k \hat{n}(0,k,0) e^{\frac{1}{2}vk^2 - \frac{1}{2}\lambda^2 k^2} H_m(k\lambda) \tag{24}$$

Under the scenario where the initial population is a delta function (for example), $\hat{n}(0,k,0) = 1$ and the integral can be performed, obtaining (for the even values):

$$c_{2m} = \frac{N_0 \lambda}{\sqrt{\pi} 2^{2m} 2m!} (-1)^m 2^{2m+\frac{1}{2}} \Gamma\left(m + \frac{1}{2}\right) \sqrt{\frac{1}{\lambda^2 - v}} \left(\frac{v + \lambda^2}{v - \lambda^2}\right)^m \tag{25}$$

where $N_0$ is the total number of individuals initially.

Some simplifications can be made to our solution:

$$\hat{n}(0,k,t) = e^{rt - bvt} e^{\left(\int_0^t \mathrm{d}t' (-aN(t'))\right)} \sum_m c_{2m} e^{-b\left((4m+1)\lambda^2\right)t} e^{-\frac{1}{2}k^2\left(\lambda^2 + v\right)} H_{2m}(k\lambda) \tag{26}$$

The total number of individuals is given by this equation with $k = 0$. The Hermite polynomial of rank $2m$ at $k = 0$ is equal to $\frac{(-1)^m (2m)!}{m!}$. We can substitute our expression for the value of $c_{2m}$ into the expression as well:

$$\hat{n}(0,0,t) = e^{rt - bvt} e^{\left(\int_0^t \mathrm{d}t' (-aN(t'))\right)} \sum_m c_{2m} e^{-b\left((4m+1)\lambda^2\right)t} \frac{(-1)^m (2m)!}{m!} \tag{27}$$

Fortunately, the sum has an analytical solution, yielding:

$$\hat{n}(0,0,t) = e^{rt - bvt} e^{\left(\int_0^t \mathrm{d}t' (-aN(t'))\right)} \frac{\sqrt{2} \lambda N_0 \sqrt{\frac{1}{\lambda^2 - v}} e^{-b\lambda^2 t}}{\sqrt{\frac{(\lambda^2 + v) e^{-4b\lambda^2 t}}{\lambda^2 - v} + 1}} \tag{28}$$

From here we can close the loop by recognizing that $n(0,0,t)$ is equal to $N(t)$ (as we mentioned at the start); additionally we introduce the function:

$$B(t) = \left( e^{rt - bvt} \frac{\sqrt{2} \lambda N_0 \sqrt{\frac{1}{\lambda^2 - v}} e^{-b\lambda^2 t}}{\sqrt{\frac{(\lambda^2 + v) e^{-4b\lambda^2 t}}{\lambda^2 - v} + 1}} \right) \tag{29}$$

By taking logarithms of eq. 28 and then taking the derivative, we can derive a differential equation for $N(t)$:

$$\frac{\mathrm{d}N(t)}{\mathrm{d}t} = \frac{B'(t)}{B(t)} N(t) - aN(t)^2 \tag{30}$$

In words, the entire process described above enables us to write down a modified logistic growth equation with a time-dependent birth rate $\mathrm{d}\ln B/\mathrm{d}t$. It should be noted that this function $B(t)$ will be dependent on the initial conditions specified in the process, leading to non-universal forms. Once $N(t)$ is solved, we can return to the solution in order to derive how the distribution itself (in Fourier space) is changing in time.

Therefore, the generic time evolution of a system with the harmonic death rate is given by:

$$N(t) = \frac{N_0 \exp\left(\int_0^t G(t')\mathrm{d}t'\right)}{N_0 \int_0^t a \exp\left(\int_0^{t'} G(t'')dt''\right)\mathrm{d}t' + 1} \tag{31}$$

where in this example:

$$G(t) = \frac{B'(t)}{B(t)} = r + \frac{b\left(v^2 - \lambda^4\right)\left(e^{4b\lambda^2 t} - 1\right)}{\left(\lambda^2 - v\right)e^{4b\lambda^2 t} + \lambda^2 + v} \tag{32}$$

This is the landscape-dependent growth rate. In the limit that $t \to \infty$ this expression becomes a constant, which is a good proxy to the total population as at this point the total births and deaths must cancel:

$$G_\infty = r - b(v + \lambda^2) \tag{33}$$

The total population will be this expression divided by $a$. As $b$ is a constant greater than zero, we can see here that it's preferable to have $v$ be as negative as possible (sign convention) and $\lambda$ as small as possible. Reintroducing $\lambda$:

$$G_\infty = r - b\left(v + \sqrt{\frac{\mathcal{D}}{b} + v^2}\right) \tag{34}$$

Despite the fact that technically $G_\infty$ does increase for stronger drifts, there are diminishing returns to $v$ becoming increasingly more negative due to the presence of the term in the square root.

### IV. STATIONARY DISTRIBUTION

Having handled the non-linearity, we can return to how we expect the distributions to appear. We can recall our definition of the distribution:

$$\hat{n}(0,k,t) = e^{rt - bvt} e^{\left(\int_0^t \mathrm{d}t'(-aN(t'))\right)} \sum_m c_{2m} e^{-b\left((4m+1)\lambda^2\right)t} e^{-\frac{1}{2}k^2\left(\lambda^2 + v\right)} H_{2m}\left(k\lambda\right) \tag{35}$$

For a given $N(t)$, this is the distribution in Fourier space. We can observe that there are many different modes relaxing independently. It can also be seen that the higher modes relax down to 0 much faster than the 0 mode, as the relaxation time for the mode is decreasing in the mode number. In this case, it would appear that a good approximation as to the form of distribution in $k$ space as $t \to \infty$ will be given by:

$$\hat{n}(0, k, t \to \infty) \sim \exp\left(-\frac{1}{2}k^2(\lambda^2 + v)\right) \tag{36}$$

Therefore, in real space the distribution will have the following form:

$$\hat{n}(0, \chi, t \to \infty) \sim \frac{1}{\sqrt{\lambda^2 + v}} e^{-\frac{\chi^2}{2(\lambda^2+v)}} \tag{37}$$

where we can see that negative $v$ serves to make the distribution narrower and that larger $\mathcal{D}$ broadens the distribution.

### V. ADAPTATION TO A SUDDEN CHANGE IN THE ENVIRONMENT

In order now to analyze how the adaptation to a sudden change in the environment can proceed, we can imagine the following scenario:

- an original population is left to relax in an environment $b(\chi - \chi_0)^2$. This will form an distribution with the Gaussian character shown in the previous section
- at time $t = 0$, we shift the environment to be $b\chi^2$. We can also shift the magnitude of the death rate by changing $a$.
- we observe how the population relaxed to the previous environment adapts to the new environment as a function of the magnitude of the shift.

Using the results of the previous section, we begin with a population of distribution:

$$n(\chi, t) = \frac{G_\infty}{a_{old}\sqrt{2\pi}\sqrt{\lambda^2 + v}} e^{-\frac{\chi^2}{2(\lambda^2+v)}} \tag{38}$$

This equation is the stationary (equilibrium) state of the original equation, as can be checked by substitution. We can make this equation correspond to a different landscape simply by shifting the $\chi$ variable to $\chi - \chi_0$. Taking the Fourier transform of this equation yields:

$$\hat{n}(0, k, 0) = \frac{G_\infty}{a_{old}} \exp\left(-\frac{1}{2}(v + \lambda^2)k^2\right) \exp\left(ik\chi_0\right) \tag{39}$$

Recalling our definition of the Hermite series coefficients:

$$c_m = \frac{\lambda}{\sqrt{\pi}2^m m!}\int \mathrm{d}k\hat{n}(0,k,0)e^{\frac{1}{2}vk^2-\frac{1}{2}\lambda^2k^2}H_m(k\lambda) \tag{40}$$

where the generic function $B(t)$ in terms of these functions is given by:

$$B(t) = e^{rt-bvt}\sum_{m=0}^{\infty} c_{2m}e^{-b\left((4m+1)\lambda^2\right)t}\frac{(-1)^m(2m)!}{(m)!} \tag{41}$$

Recall that the dynamics of the population subsequent to the perturbation is given by Eq. 31. The integral over the Hermite polynomials can be performed, leading to:

$$c_m = \frac{G_\infty}{a_{old}2^m m!}\frac{\chi_0^m e^{-\frac{\chi_0^2}{4\lambda^2}}}{(-i)^m\lambda^m} \tag{42}$$

Inputting this expression into the formula for $B(t)$ again yields an analytic expression after the summation is performed:

$$B(t) = e^{rt-bvt}\frac{\sqrt{\pi}G_\infty}{a_{old}}e^{-\frac{4b\lambda^4t+\chi_0^2}{4\lambda^2}}\cosh\left(\frac{x_0e^{-2b\lambda^2t}}{2\lambda}\right) \tag{43}$$

This leads to the growth function $G(t)$ of the form:

$$G(t) = r - b\left(\lambda^2+v\right) - b\lambda x_0 e^{-2b\lambda^2 t}\tanh\left(\frac{x_0e^{-2b\lambda^2t}}{2\lambda}\right) \tag{44}$$

where it is easy to observe that over long timescales this growth function returns back to our previous form. However, we are now interested in the dynamics of adaption to this new environment. To that end we need to take the integral of $G(t)$ between 0 and $t$.

$$\int_0^t \mathrm{d}t'G(t') = -bt\left(\lambda^2+v\right) + \log\left(\cosh\left(\frac{x_0e^{-2b\lambda^2t}}{2\lambda}\right)\right) + rt - \log\left(\cosh\left(\frac{x_0}{2\lambda}\right)\right) \tag{45}$$

Although the expression in the denominator of $N(t)$ has no analytic form, it can be calculated numerically. This again shows that populations with large $v$ adapt better to a sudden change. Plots of various scenarios are displayed over the next few pages.

The relaxation time in the main text is calculated by applying a shift $\chi_0$ to a stationary population and measuring how long it takes for the population to recover to 99% of its equilibrium value.

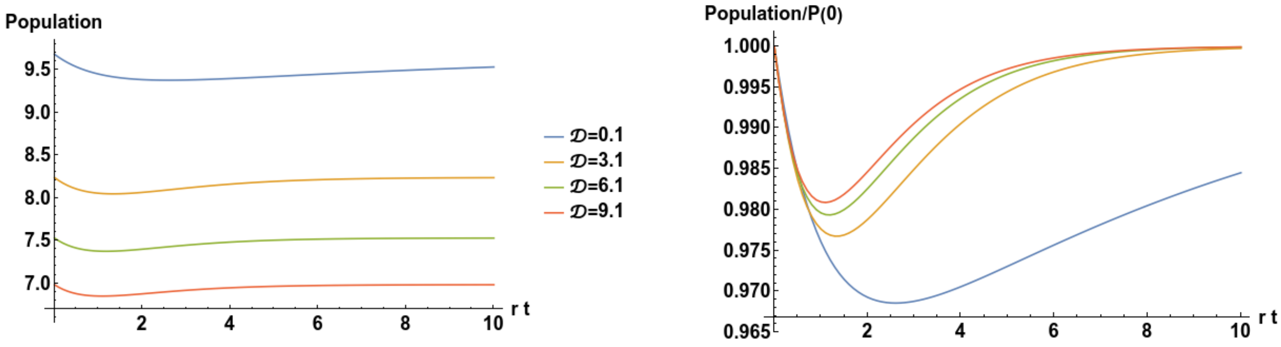


FIG. 2. Recovery of a population with $v = 0$ to a perturbation to $\chi$ for different $D$. Populations with a larger $D$ have a smaller initial population, but a faster recovery time. (a=1,b=1)

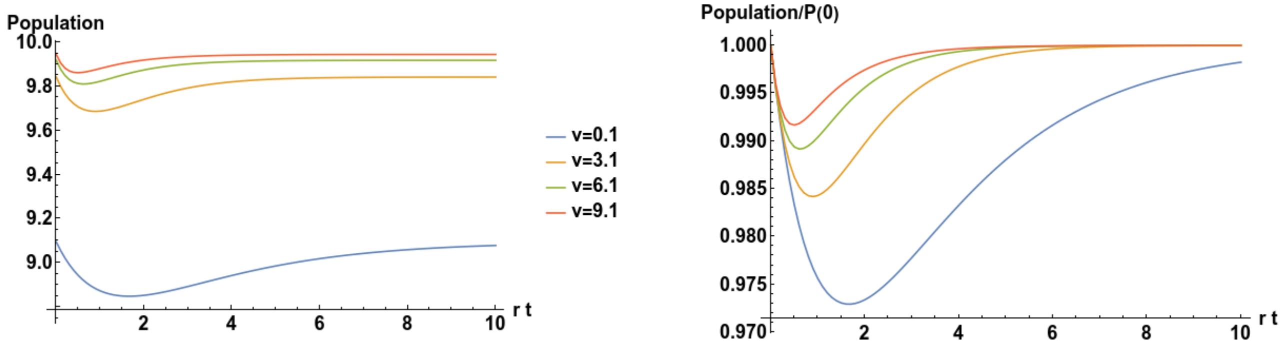


FIG. 3. Recovery of a population with $D = 1$ to a perturbation to $\chi$ for different values of $v$. Unlike the prior figure, populations with larger $v$ have a larger initial population, but also a faster recovery time.(a=1,b=1)

### VI. CONTINUAL VARIATION OF THE ENVIRONMENT

In order to analyze the effect of fast environmental switching, we shall first have to expound the nature of our solution under an arbitrary initial condition. One way that the equations so far can simulate dynamics is as follows:

- Select a starting initial condition $\hat{n}(0, k, 0)$ under some death probability
- Solve for

$$c_m = \frac{\lambda}{\sqrt{\pi} 2^m m!} \int \mathrm{d}k \hat{n}(0, k, 0) e^{\frac{1}{2} v k^2 - \frac{1}{2} \lambda^2 k^2} H_m (k\lambda)$$

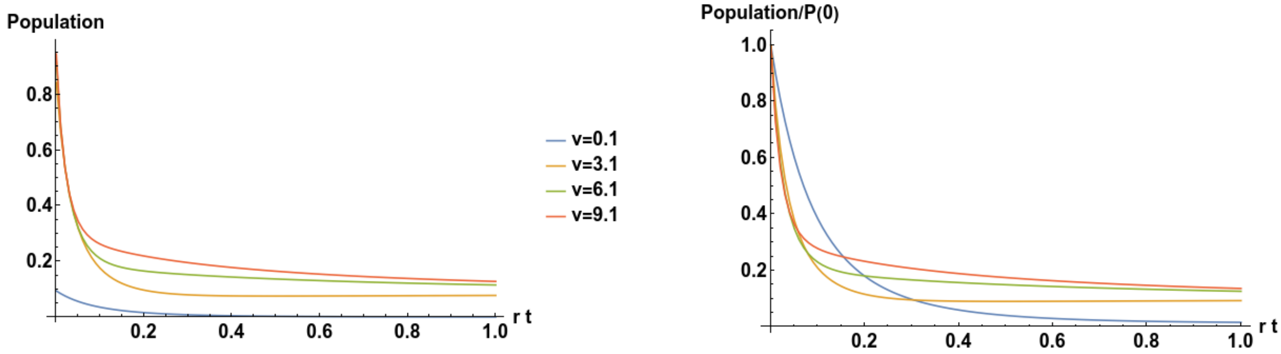


FIG. 4. Recovery of a population with $D = 1$ to a perturbation to $\chi$ and an additional increase in the death rate due to the change of environment, from $a = 1$ to $a = 10$, for different values of $v$. Populations with larger $v$ have a larger initial population, but the effects of an environmental change such as this (reduction in niche size) are more catastrophic for the fast adaptors than the slow adaptors. (b=1)

- Solve for

$$B(t) = \sum_m c_{2m} e^{-b\left((4m+1)\lambda^2\right)t} \frac{(-1)^m (2m)!}{m!}$$

- Solve for

$$N(t) = \frac{e^{rt-bvt} B(t) N(0)}{B(0) + aN(0) \int_0^t \mathrm{d}t' e^{rt'-bvt'} B(t')}$$

- Solve for the final distribution

$$\hat{n}(0,k,t) = e^{rt-bvt} e^{\left(\int_0^t \mathrm{d}t' (-aN(t'))\right)} \sum_m c_m e^{-b\left((2m+1)\lambda^2\right)t} e^{-\frac{1}{2}k^2\left(\lambda^2+v\right)} H_m\left(k\lambda\right)$$

- Shift the distribution by multiplying by $e^{ik\chi_0}$ and then use that distribution as an input and repeat

We will here distinguish between two types of shift strategies. When we multiply the distribution by $e^{ik\chi_0}$, we are in fact establishing that the population prior to the shift in the death probability was relaxing to the death rate $(\chi - \chi_0)^2$. We can either choose that the $\chi_0$ we shift to are alternating between the same two points, hence allowing for lack of adaption to be good once the system returns back to its "initial" environment. Alternatively, we can choose $\chi_0$ to be constantly moving in the same direction, such that the system is trying to relax to a completely new environment each time. In this case, lack of adaption is even more detrimental. We might anticipate that these two scenarios entail different dynamics.

### VII. CONDITIONS ON SURVIVAL

In order to derive conditions on survival and extinction of populations, we consider the behavior of populations when subjected to a single shift. From this, we can infer the properties of repeated applications of environmental shifts on the population.

We begin with the time evolution equation of each mode $c_m(t)$, which is given by:

$$\frac{\mathrm{d}c_m(t)}{\mathrm{d}t} = (N_{eq}a - 2mb\lambda^2 - aN(t))c_m(t) \tag{46}$$

In order to estimate relaxation times, we focus on the zeroth mode of the system, which is also the largest mode:

$$\frac{\mathrm{d}c_0(t)}{\mathrm{d}t} = (N_{eq}a - aN(t))c_0(t) \tag{47}$$

Our estimate of the timescale shall proceed from the following facts: at equilibrium, only the zeroth mode of the system is non-zero, $c_m = 0, m > 0$, and we want the population to be the same, $N(\Delta t) = N(0)$. This corresponds to full relaxation of the zeroth mode after a shift. Equation (47) is solved by:

$$c_0(\Delta t) = c_0(0) \exp\left(N_{eq}a\Delta t - a\int_0^{\Delta t} N(t)\mathrm{d}t\right) \tag{48}$$

The value $c_0(0)$ can be calculated by considering the equilibrium state and applying a shift of $\Delta\chi$ to it, then recalculating the value of $c_0(0)$. This is given by:

$$c_0(0) = N(0)\exp(-\Delta\chi^2/4\lambda^2) \tag{49}$$

We can also use the above expression to see that the relaxation to the new equilibrium is complete when when $c_0(\Delta t) = N(0)$. We can therefore rearrange equation (49) to yield the following condition:

$$N(0) = N(0)\exp(-\Delta\chi^2/4\lambda^2)\left(N_{eq}a\Delta t - a\int_0^{\Delta t} N(t)\mathrm{d}t\right) \tag{50}$$

where $N(0)$, the initial population number, cancels out. Equation (50) defines the condition for full relaxation of the first mode after a shift in the environment is applied. As $aN(t)$ is always positive, we can derive a rather simple condition from equation (50), which is that, minimally:

$$-\Delta\chi^2/4\lambda^2 + N_{eq}a\Delta t > 0 \tag{51}$$

in order for full relaxation of the zeroth mode to be completed in a time $\Delta t$ after a shift $\Delta\chi$

### VIII. FERTILITY, INHERITANCE AND THEIR EFFECT ON ADAPTATION TO ENVIRONMENT

We proceed to extend our baseline model to more complicated scenarios by relaxing both the $I = \delta(\chi - \chi')$ condition, as well as the $R = 0$ assumption. The first order effect of imperfect inheritance can be captured through resetting of the diffusion coefficient, as we shall demonstrate for a Gaussian inheritance function

$$I(\chi - \chi') = (1/\sqrt{2\pi\sigma^2})\exp(-(\chi - \chi')^2/(2\sigma^2)). \tag{52}$$

We will use the convolution theorem to Fourier transform the first term in the second of Eqs. 1 into $k$-space:

$$\int_{-\infty}^{\infty} \mathrm{d}\chi \exp(-ik\chi) \int P_R(T')I(\chi - \chi')n(T', \chi', t)\mathrm{d}\chi'\mathrm{d}T' = r\exp\left(-\frac{1}{2}k^2\sigma^2\right)\hat{n}(R, k, t) \tag{53}$$

Taylor expansion of the exponential yields the following:

$$r\left(1 - \frac{1}{2}k^2\sigma^2 + \frac{1}{4}k^4\sigma^4 - \dots\right)\hat{n}(R, k, t) \tag{54}$$

For a not too large value of $\sigma$, we ignore the corrections going as $\mathcal{O}(k^4)$ and higher. This might yield inaccurate solutions at large values of $k$, however, given how we designed the death rate, the number of individuals at large values of $k$ is very small. To first order then we can see that the effect of imperfect inheritance can be given simply by:

$$D \to D + \frac{1}{2}r\sigma^2 \tag{55}$$

The other extension, accounting for declining fertility ($R > 0$) means that our transformed equations now exist in two modes, 0 and $R$:

$$\frac{\partial\hat{n}(R, k, t)}{\partial t} = (r - R)\hat{n}(R, k, t) + 2vbk\frac{\partial\hat{n}(R, k, t)}{\partial k} - Dk^2\hat{n}(R, k, t) - a\hat{n}(R, k, t)N(t) + b\frac{\partial^2\hat{n}(R, k, t)}{\partial k^2} \tag{56}$$

$$\frac{\partial\hat{n}(0, k, t)}{\partial t} = r\hat{n}(R, k, t) + 2vbk\frac{\partial\hat{n}(0, k, t)}{\partial k} - Dk^2\hat{n}(0, k, t) - a\hat{n}(0, k, t)N(t) + b\frac{\partial^2\hat{n}(0, k, t)}{\partial k^2} \tag{57}$$

This set of equations appears decidedly more complex than the previous ones, however, they have a similar stationary state, which is given by:

$$\hat{n}_S(0,k) = \frac{1}{a}\left(r - R - b\left(v + \sqrt{\frac{\mathcal{D}}{b} + v^2}\right)\right)\exp\left(-\frac{1}{2}\left(v + \sqrt{\frac{D}{b} + v^2}\right)k^2\right) \tag{58}$$

$$\hat{n}_S(R,k) = \frac{r-R}{ra}\left(r - R - b\left(v + \sqrt{\frac{\mathcal{D}}{b} + v^2}\right)\right)\exp\left(-\frac{1}{2}\left(v + \sqrt{\frac{D}{b} + v^2}\right)k^2\right) \tag{59}$$

The mode $R$, which tells us how much the species is reproducing, is equal to the mode 0 multiplied by the factor $(r-R)/r$. Moreover, it is possible to prove that once the system has reached a point where $n(R,k,t) = ((r-R)/r)n(0,k,t)$ no further dynamics in the ratio of $n(R,k,t)/n(0,k,t)$ occur, no matter the form of environmental change. This means that the effective dynamics reduce again down to a single mode even with declining fertility. We can prove this by analyzing the following derivative:

$$\frac{\partial}{\partial t}\left(\frac{\hat{n}(R,k,t)}{\hat{n}(0,k,t)}\right) = \frac{\hat{n}(0,k,t)\hat{n}'(R,k,t) - \hat{n}(R,k,t)\hat{n}'(0,k,t)}{\hat{n}(0,k,t)^2} \tag{60}$$

where the equation has been derived using the quotient rule ($n'$ refers here to derivative with respect to t). The differential equations given in 56 and 57 can be substituted into equation 60.

We can rewrite equations 56 and 57 as:

$$\frac{\partial \hat{n}(R,k,t)}{\partial t} = \hat{O}\hat{n}(R,k,t) + (r-R)\hat{n}(R,k,t) \tag{61}$$

$$\frac{\partial \hat{n}(0,k,t)}{\partial t} = \hat{O}\hat{n}(0,k,t) + r\hat{n}(R,k,t) \tag{62}$$

where $\hat{O}$ is a linear operator. Substituting into 60 yields:

$$\frac{\partial}{\partial t}\left(\frac{\hat{n}(R,k,t)}{\hat{n}(0,k,t)}\right) = \frac{\hat{n}(0,k,t)(\hat{O}\hat{n}(R,k,t) + (r-R)\hat{n}(R,k,t)) - \hat{n}(R,k,t)(\hat{O}\hat{n}(0,k,t) + r\hat{n}(R,k,t))}{\hat{n}(0,k,t)^2} \tag{63}$$

Now we imagine that one of the modes is given as a multiple of another, for example $\hat{n}(R,k,t) = c\hat{n}(0,k,t)$. We could, for instance, instantiate the system in this configuration if we wished. This expression can be cleaned up, yielding:

$$\frac{\partial}{\partial t}\left(\frac{\hat{n}(R,k,t)}{\hat{n}(0,k,t)}\right) = c(r-R) - rc^2 \tag{64}$$

We make the observation that the dynamics of the quotient between modes $R$ and 0 depends only on a constant if the two modes are initially a multiplicative constant of one another.

When $c = (r-R)/r$, the right hand side of expression 60 is zero, i.e. the modes no longer change their relative magnitude compared to one another.

This proof doesn't demonstrate that the system will always be found in this state, but it does show that under a special condition we can ignore the dynamics of one of the modes. Moreover, analysis of equations 56 and 57 shows that both equations have the same dependence on $k$. In other words, they should approach the same distribution in $k$ as time increases, but, as we already demonstrated, if the two modes are merely multiples of one another (which occurs when they have the same distribution in $k$) we can dispense with one of the modes. Therefore the model with declining fertility, despite appearing more complicated, can be mapped onto a single mode equation through the substitution $\hat{n}(R,k,t) \to ((r-R)/r)\hat{n}(0,k,t)$ and we can be assured nothing funny will occur in the other modes.

Taking as our key mode $S = 0$, we have the following equation for a system with declining fertility and imperfect inheritance:

$$\frac{\partial \hat{n}(0,k,t)}{\partial t} = (r-R)\hat{n}(0,k,t) + 2vbk\frac{\partial \hat{n}(0,k,t)}{\partial k} - (D + \frac{1}{2}r\sigma^2)k^2\hat{n}(0,k,t) - a\hat{n}(0,k,t)N(t) + b\frac{\partial^2 \hat{n}(0,k,t)}{\partial k^2} \tag{65}$$

whereupon we see that if we define $\tilde{r} = r - R$ and $\tilde{D} = (D + \frac{1}{2}r\sigma^2)$ we recover the perfect inheritance, no declining fertility model with new values of the replication and diffusion.

To realize the full potential of this model, we need to non-dimensionalize it. To this end we imagine the following substitutions:

$$\hat{n}(0,k,t) = \tilde{N}\tilde{n}(0,k,t) \tag{66}$$

$$k = k_c\tilde{k} \tag{67}$$

$$t = t_c\tau \tag{68}$$

$$\tilde{N} = \frac{r-R}{a} \tag{69}$$

This yields the following:

$$\begin{aligned}\frac{N}{t_c}\frac{\partial \tilde{n}(0,k,\tau)}{\partial \tau} =& (r-R)\tilde{N}\tilde{n}(0,k,\tau) + 2vb\tilde{N}\tilde{k}\frac{\partial \tilde{n}(0,\tilde{k},\tau)}{\partial \tilde{k}} - \tilde{N}k_c^2(D + \frac{1}{2}r\sigma^2)\tilde{k}^2\tilde{n}(0,\tilde{k},\tau) \\ & - a\tilde{N}^2\tilde{n}(0,\tilde{k},\tau)\tilde{n}(0,0,\tau) + \frac{b\tilde{N}}{k_c^2}\frac{\partial^2 \tilde{n}(0,\tilde{k},\tau)}{\partial \tilde{k}^2}\end{aligned} \tag{70}$$

dividing through by $\tilde{N}/t_c$ gives:

$$\begin{aligned}\frac{\partial\tilde{n}(0,k,\tau)}{\partial\tau} =&(r-R)t_c\tilde{n}(0,k,\tau)+2vb\tilde{k}\frac{\partial\tilde{n}(0,\tilde{k},\tau)}{\partial\tilde{k}}-t_ck_c^2(D+\frac{1}{2}r\sigma^2)\tilde{k}^2\tilde{n}(0,\tilde{k},\tau) \\ &-aNt_c\tilde{n}(0,\tilde{k},\tau)\tilde{n}(0,0,\tau)+\frac{bt_c}{k_c^2}\frac{\partial^2\tilde{n}(0,\tilde{k},\tau)}{\partial\tilde{k}^2}\end{aligned} \tag{71}$$

where:

$$t_c = 1/(r-R) \tag{72}$$

$$k_c^2 = \frac{r-R}{D+\frac{1}{2}k\sigma^2} \tag{73}$$

This equation reduces to:

$$\begin{aligned}\frac{\partial\tilde{n}(0,k,\tau)}{\partial\tau} =&\tilde{n}(0,k,\tau)+\frac{2vb}{r-R}\tilde{k}\frac{\partial\tilde{n}(0,\tilde{k},\tau)}{\partial\tilde{k}}-\tilde{k}^2\tilde{n}(0,\tilde{k},\tau) \\ &-\tilde{n}(0,\tilde{k},\tau)\tilde{n}(0,0,\tau)+\frac{b(D+\frac{1}{2}r\sigma^2)}{(r-R)^2}\frac{\partial^2\tilde{n}(0,\tilde{k},\tau)}{\partial\tilde{k}^2}\end{aligned} \tag{74}$$

where there are only two dimensionless variables affecting the temporal change of the reduced number of individuals in a population:

$$\tilde{v} = \frac{vb}{r-R} \text{ and } \tilde{D} = \frac{b(D+\frac{1}{2}r\sigma^2)}{(r-R)^2} \tag{75}$$

We conclude that, when measured in units of population of $(r-R)/a$ and units of time of $1/(r-R)$, two systems with the same value of $\tilde{v}$ and of $\tilde{D}$ will have exactly the same behavior.

### IX. $\chi$-DEPENDENT REPLICATION

In the previous sections we considered the case in which the probability of death per unit time is affected by some state variable $\chi$. We could imagine a different form of our model, one in which the probability to replicate depends on this variable instead. We may ask whether this model is isomorphic to the death rate model. To proceed, we write our equations in this updated form. For simplicity, we shall take the perfect inheritance version of our model $I(\chi-\chi') = \delta(\chi-\chi')$

We therefore have the following set of equations:

$$\frac{\partial n\left(T,\chi,t\right)}{\partial t} = -\frac{\partial n\left(T,\chi,t\right)}{\partial T} - \frac{\partial}{\partial \chi}\left(f(\chi)n(T,\chi,t)\right) + D\frac{\partial^2 n(T,\chi,t)}{\partial \chi^2} - P_D(T,\chi)n(T,\chi,t) \tag{76}$$

$$n(0,\chi,t) = \int P_R(T',\chi)n(T',\chi,t)\mathrm{d}T' \tag{77}$$

$$n(\infty,x,t) = 0 \tag{78}$$

Another assumption that we make is that the replication function $P_R$ is separable, $P_R(\chi,T) = \mathcal{X}(\chi)\mathcal{T}(T)$. The age-dependent effective replication rate will be given by:

$$\mathcal{T}(T) = r\exp(-RT) \tag{79}$$

We can take the age-related Laplace transform once more:

$$\frac{\partial \hat{n}\left(S,\chi,t\right)}{\partial t} = -S\hat{n}\left(S,\chi,t\right)+r\mathcal{X}(\chi)\hat{n}\left(R,\chi,t\right)-\frac{\partial}{\partial \chi}\left(f(\chi)\hat{n}(S,\chi,t)\right)+D\frac{\partial^2 \hat{n}(S,\chi,t)}{\partial \chi^2}-P_D(T,\chi)\hat{n}(S,\chi,t) \tag{80}$$

Let's consider the $R = 0$ case, leading to a model with a single mode:

$$\frac{\partial \hat{n}\left(0,\chi,t\right)}{\partial t} = r\mathcal{X}(\chi)\hat{n}\left(0,\chi,t\right)-\frac{\partial}{\partial \chi}\left(f(\chi)\hat{n}(0,\chi,t)\right)+D\frac{\partial^2 \hat{n}(0,\chi,t)}{\partial \chi^2}-P_D(T,\chi)\hat{n}(0,\chi,t) \tag{81}$$

This model is not identical to our baseline model. For one, we can see that the function $\mathcal{X}(\chi)$ must have a different form to the death rate probability we had previously. For if it were to go as $\sim \chi^2$ there would be regions where the probability to replicate would become very large as $|\chi|$ increases (N.B. this isn't a problem for the death rate). Similarly, a function of the form $-\chi^2$ would have the issue that the replication rate would become negative at certain points. Therefore, this function would have to be bounded to be realistic. It is not possible to straightforwardly map the results of our analytical model to this system.

### X. MORE REALISTIC BIRTH RATES

We introduced a form of birth rate which is obviously inaccurate, that is that the birth rate probability is given by:

$$P_R(T) = r\exp(-RT) \tag{82}$$

which has the desired feature that fertility declines with age, but assumes unrealistically, that maximal fertility occurs at the moment of birth. We wish to conceive of a different scheme where fertility declines at large ages, but also starts at zero, meaning that organisms have some period of time before they begin to reproduce. In order to be able to preserve Laplace transforms for analytical simplicity, we introduce the following form of $P_R(T)$ that has the desired features:

$$P_R(T) = r\left(\exp(-R_1 T) - \exp(-R_2 T)\right) \tag{83}$$

So long as $R_2 > R_1$, we have a function that starts at zero at $T = 0$, and decays to zero as $T \to \infty$, but with finite reproduction at intermediate ages, with a maximum at $(\ln R_1/R_2)/(R_1 - R_2)$

We reuse the operator introduced in equations (61) to succinctly write down the time evolution equations for this more complicated model. In such a system we only have 3 important modes:

$$\frac{\partial \hat{n}(0,k,t)}{\partial t} = r(\hat{n}(R_1,k,t) - \hat{n}(R_2,k,t)) + \hat{O}\hat{n}(0,k,t) \tag{84}$$

$$\frac{\partial \hat{n}(R_1,k,t)}{\partial t} = (r - R_1)\hat{n}(R_1,k,t) - r\hat{n}(R_2,k,t) + \hat{O}\hat{n}(R_1,k,t) \tag{85}$$

$$\frac{\partial \hat{n}(R_2,k,t)}{\partial t} = r\hat{n}(R_1,k,t) - (r + R_2)\hat{n}(R_2,k,t) + \hat{O}\hat{n}(R_2,k,t) \tag{86}$$

Now we follow the familiar procedure of looking at how the ratios of these modes to one another appear. As, once again, the evolution in $k$-space is identical for every mode, these parts can be separated out and we can analyze the ratios of these modes to one another, i.e., we search for solutions where $\hat{n}(R_1,k,t) = c_1\hat{n}(0,k,t)$ and $\hat{n}(R_2,k,t) = c_2\hat{n}(0,k,t)$, we can employ the quotient rule and the equations (84)-(86) for this purpose:

$$\frac{\partial}{\partial t}\left(\frac{\hat{n}(R_1,k,t)}{\hat{n}(0,k,t)}\right) = (c_1 - 1)(c_1 - c_2)(-r) - c_1 R_1 \tag{87}$$

$$\frac{\partial}{\partial t}\left(\frac{\hat{n}(R_2,k,t)}{\hat{n}(0,k,t)}\right) = (c_2 - 1)(c_2 - c_1)\, r - c_2 R_2 \tag{88}$$

Once again, this suggests that there are particular values of the numbers $c_1$ and $c_2$ in which the quotients of the modes do not display any time evolution. In other words, the apparent fact that we have three modes can again be reduced to a single mode, despite the additional complexity. The equations above do in fact have the following solutions:

$$\begin{array}{ll} c_1 = 0 & c_2 = 0 \\ c_1 = -\frac{R_1\left(R_1-R_2+\sqrt{(R_1-R_2-4)(R_1-R_2)}-2\right)+2R_2}{2(R_1-R_2)} & c_2 = \frac{1}{2}\left(-\frac{\sqrt{(R_1-R_2-4)(R_1-R_2)}R_2}{R_1-R_2}+R_2+2\right) \\ c_1 = \frac{R_1\left(-R_1+R_2+\sqrt{(R_1-R_2-4)(R_1-R_2)}+2\right)-2R_2}{2(R_1-R_2)} & c_2 = \frac{1}{2}\left(\frac{\sqrt{(R_1-R_2-4)(R_1-R_2)}R_2}{R_1-R_2}+R_2+2\right). \end{array} \tag{89}$$

Of these solutions, only the second displays the property that $c_1 > c_2$ when $R_2 > R_1 > 0$, which is a relevant feature for real distributions as we expect the number of individuals to decline with age (we expect $\int_0^\infty \mathrm{d}T n(T,x,t)\exp(-RT)$ to decrease as $R$ increases).

Inserting these values into the equation for $\hat{n}(0,k,t)$ and simplifying leads to the following equation:

$$\frac{\partial \hat{n}(0,k,t)}{\partial t} = \left(\frac{1}{2}\left(\sqrt{(R_2-R_1)(4r-R_1+R_2)}-R_1-R_2\right)\right)\hat{n}(0,k,t)+2vbk\frac{\partial \hat{n}(0,k,t)}{\partial k}- \tag{90}$$

$$Dk^2\hat{n}(0,k,t)-a\hat{n}(0,k,t)N(t)+b\frac{\partial^2\hat{n}(0,k,t)}{\partial k^2}$$

which, as we can see, is identical to the equations we have dealt with in the main text. The only remaining factor is to determine whether the fixed point is one to which the system naturally ends up in, which can be studied by ensuring that the flow of equations (87) and (88) is always towards the fixed point, even if it wasn't initialized at the fixed point. This can be checked graphically by looking at the flow plot against $c_1, c_2$ for a given value of $r, R_1, R_2$. These graphs display a flow towards the fixed point value so long as there is a non-zero population that exists at this value. When considering long time phenomena, one can ignore initial transients associated with non-equilibrated values of $c_1, c_2$ and just observe what were to occur in such systems for the reduced model.
